# Interactions between sensory-biased and supramodal working memory networks in the human cerebral cortex

**DOI:** 10.1101/2025.07.17.665351

**Authors:** Thomas Possidente, Vaibhav Tripathi, Joseph T. McGuire, David C. Somers

## Abstract

Human working memory is supported by a broadly distributed set of brain networks. Content-specific networks communicate with a domain-general, supramodal network that is recruited regardless of the type of content. Here, we contrasted visual and auditory working memory tasks to examine interactions between the supramodal network and two content-specific networks. Functional connectivity among visual-biased, auditory-biased, and supramodal working memory networks was assayed by collecting task and resting-state fMRI data from 24 human participants (age 18-43; 11 men and 13 women). At rest, as found previously, the supramodal network exhibited stronger functional connectivity with the visual-biased network than with the auditory-biased network. This asymmetry raises questions about how networks communicate to support robust performance across modalities. However, during auditory task performance, dynamic changes increased auditory network connectivity with supramodal and visual-biased frontal regions, while decreasing connectivity from posterior visual areas to supramodal and frontal visual regions. In contrast, the visual task produced weak changes. Across individuals, auditory working memory precision correlated with the strength of auditory network connectivity changes, while no such brain-behavior link was observed for visual working memory. These results demonstrate an asymmetry in working memory network organization and reveal that dynamic reorganization accompanies performance of working memory tasks.

## Introduction

Human working memory (WM) is supported both by ‘content-specific’ mechanisms that vary with the type of information held in memory and by ‘multiple-demand’ or ‘central executive’ mechanisms whose operation appears independent of content (e.g., Baddeley & Hitch, 1974; Duncan & Owen, 2000). The interaction between these subsystems is an important, understudied aspect of WM. Sensory WM for visual stimuli and auditory stimuli has been shown to recruit different modality-specific structures in occipito-parietal and temporal lobes, respectively (Bettencourt & Xu, 2016; Christophel et al., 2012, 2017; Ester et al., 2015; Harrison & Tong, 2009; Linke et al., 2011; Serences et al., 2009; Uluç et al., 2018), while both modalities recruit broad regions of lateral frontal cortex which could be considered ‘supramodal’ or ‘multiple-demand’ in their functionality (Assem et al., 2020; Badre & Nee, 2018; Christophel et al., 2017; Crittenden & Duncan, 2014; D’Esposito & Postle, 2015; Duncan & Owen, 2000) Additionally, multiple regions within lateral frontal cortex are biased toward either visual or auditory WM (Assem et al., 2022; Mayer et al., 2017; Michalka et al., 2015; Noyce et al., 2017, 2022; Tobyne et al., 2017). These small regions were revealed by precision mapping of lateral frontal cortex in individual subjects whereas prior group-averaged analyses had blurred these regions together and mischaracterized them as ‘multiple-demand’. Moreover, these sensory-biased regions lie in proximity to true supramodal or multiple-demand WM regions (Assem et al., 2022). This motivates new questions about how these sensory-biased and domain-general areas are topographically organized and how they might functionally interact. Here, we examine how sensory-biased and supramodal WM regions interact during visual WM and auditory WM tasks and during rest.

Assem et al. (2022) suggests that sensory-biased regions surround and potentially feed information into the domain-general areas, while Noyce et al. (2017) indicates that visual-biased areas, but not auditory-biased areas, contain some domain-general response properties while still being functionally separable from fully domain-general areas. This and other work from our laboratory (Noyce et al., 2022; Tobyne et al., 2017, 2025) points to a potential functional asymmetry between visual and auditory WM mechanisms; visual cognition structures in frontal cortex appear more closely integrated with supramodal WM brain structures (regions that respond to both auditory and visual WM demands) than are auditory cognition structures in frontal cortex. There is also a substantial behavioral literature demonstrating visual dominance over auditory or tactile modalities and these effects appear to depend on attentional interactions (Colavita, 1974; Hartcher-O’Brien et al., 2008; Hecht & Reiner, 2009; Posner et al., 1976; Spence et al., 2012). One natural question is how it is possible for auditory WM to function effectively with relatively weaker coordination with supramodal networks. Therefore, we hypothesize that visual and auditory WM task performance evoke asymmetric changes in functional connectivity between visual-biased, auditory-biased, and supramodal networks. We employ individual-specific task-based functional connectivity analysis, which has substantially greater power than standard group analysis (e.g., Gratton et al., 2018). Several non-mutually exclusive mechanisms are plausible to help explain how auditory WM performance acquires the necessary resources to function effectively despite weaker coordination with supramodal WM networks at rest: auditory WM performance may 1) increase connectivity between auditory and supramodal networks, 2) decrease competing connectivity between visual and supramodal networks, 3) increase connectivity within the auditory network components, and/or 4) increase cross modal connectivity between visual frontal and auditory frontal networks. An increase in connectivity in the auditory network components (*Mechanism 3*) could reflect increased efficiency within the auditory system. An increase in cross-modal connectivity (*Mechanism 4*) would increase direct communication between the auditory-biased and the frontal visual-biased regions, which exhibit partial supramodal WM properties, and could indirectly increase communication with supramodal network regions based on the resting connection between visual-biased and supramodal regions.

The present study examines these possibilities by presenting in-depth resting state and task-based functional connectivity analyses of sensory-biased perception and WM regions, along with supramodal frontal regions. Building on our prior work (Michalka et al., 2015; Noyce et al., 2017, 2022; Tobyne et al., 2025), we define a large number of visual-biased, auditory-biased, and supramodal WM regions of interest (ROIs) in the frontal cortex of individual subjects to drive our analysis. We replicate the finding that the supramodal network is more strongly connected with the visual-biased network than with the auditory-biased network at rest and additionally find that visual-biased, auditory-biased, and supramodal networks change their connectivity structure differentially during visual and auditory WM tasks. We show that the auditory-biased network overcomes its resting-state connectivity disadvantage by recruiting both supramodal and visual-biased networks, increasing connectivity within the auditory network, and by effecting a decrease in competing connectivity among visual frontal, visual posterior, and supramodal networks. We also show that task-based changes in functional connectivity between a subset of these networks are predictive of behavioral WM precision, indicating that increased recruitment by the auditory network leads to increased auditory WM performance.

## Methods

### Participants

Twenty-four participants (aged 18 to 43; 11 men and 13 women [self-identified]) were recruited from the Boston University community. All procedures in this study were approved by the Institutional Review Board of Boston University (4928E). Each participant gave written informed consent to participate in the study and received monetary compensation. All participants indicated that they had normal or corrected to normal vision and normal hearing. Two authors participated in both the task and resting state scans (T. P., V. T.). All 24 subjects performed visual and auditory working memory and sensorimotor control tasks during fMRI for this study. Twenty-two of these also participated in resting-state scans. Three participants were rejected from all analyses - two for ROI identification problems, one for head movement – leaving twenty-one participants (19 with resting state scans). One additional participant was rejected from only the resting state analysis for head movement, leaving eighteen participants. Rejection criteria details are given below.

### Working Memory Paradigm

All participants performed 4 tasks in the scanner: 2-back visual working memory, 2-back auditory working memory, visual sensorimotor control, and auditory sensorimotor control. The stimuli and general paradigm are shown in Figure 1 below. Stimuli display and task timing control for the 2-back WM tasks were performed using the Psychopy Toolbox (v2021.1.4; www.psychopy.org; Pierce et al., 2019) in Python (v. 3.10 www.python.org) running on a laptop running Windows 11 and displayed using a liquid crystal display projector that back-projected onto a screen placed within the scanner bore. Auditory stimuli were presented through MRI-safe in–ear earphones (Sensimetrics Model S14; Sensimetrics; www.sens.com). Stimuli were presented for 1 second each, with a 1 second gap between stimuli; only one modality was presented at a time. The visual stimuli were Gabor patches with spatial frequencies in the range of 1 to 6 cycles per degree. In the visual 2-back task, subjects judged whether the spatial frequency of the stimulus matched that presented two stimuli previously. After the presentation of the first two stimuli in a block, matches occurred on 50% of the trials. Gabor orientation varied randomly and was irrelevant to the match judgement. In the auditory 2-back task, the stimuli were auditory ‘warbles’ with either a 500 or 1000 Hz base pitch modulated by an oscillatory amplitude envelope with frequencies ranging from 3 Hz to 20 Hz. Subjects judged whether the envelope frequency matched that from two stimuli earlier. Pitch was varied randomly and was irrelevant to the match judgement. For both auditory and visual stimuli, the trials consisted of high frequency stimuli (i.e., odd numbered trials) alternated with low frequency trials (i.e., even numbered trials); this afforded the opportunity to make the matching task difficult while also presenting a broader range of stimuli. We sought to equate task difficulty and WM load across modalities for each individual by manipulating WM precision (Alvarez & Cavanagh, 2004; Bays et al., 2009; Franconeri et al., 2013). The percentage change in the modulatory frequency for non-match trials was adjusted on a trial-by-trial basis using independent 3-up/1-down staircasing procedures for visual and auditory stimuli (stepsize of 1%). Individual thresholds were identified for each subject in a behavioral session prior to scanning and were updated during the scans. The behavioral training session included auditory presentation of recorded acoustic noise of the scanner to simulate the auditory distractions of the scanner. Individual differences in working memory precision were quantified by the staircase threshold parameters. Smaller thresholds imply more precise performance, even though task accuracy was equated across subjects. These threshold measures were employed for individual-subject measures comparing functional connectivity changes and behavioral changes.

**Figure 1.**
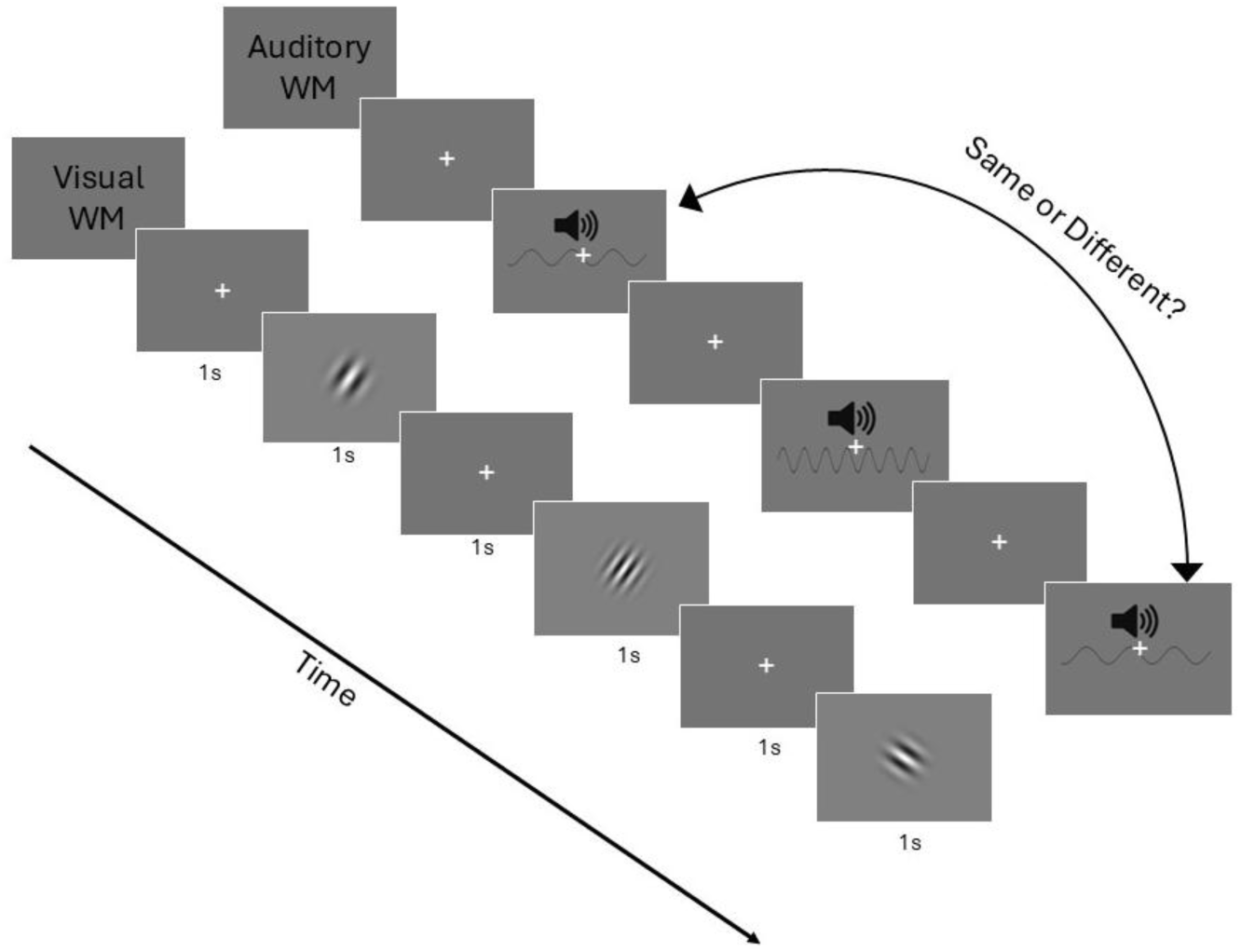
Visual and auditory 2-back WM task summaries. For the visual WM task, participants were asked whether the current Gabor filter spatial frequency matched the one presented two stimuli previously. For the auditory WM task participants were asked whether the warble envelope volume modulation frequency matched the one presented two stimuli previously. Task performance required participants to maintain a high-precision representation of both the most recent even and odd numbered trials.

During the sensorimotor control tasks, auditory or visual stimuli were presented without 2-back repeats, and participants were instructed not to do the task but instead to arbitrarily press either of the two response buttons at the end of each trial. The relevant blocks in a single run consisted of 2 blocks of visual WM, 2 blocks of auditory WM, 1 block of visual sensorimotor control, 1 block of auditory sensorimotor control, and 1 fixation condition block without stimuli or responses. Block order was randomized from run to run for all participants. Each block lasted 32s and contained 16 trials. Participants were given feedback indicating correct and incorrect trials. During the sensorimotor control conditions, feedback was given if they had pressed the key or not. For participants used in the task analyses, 19 had four runs, one had three, and one had six.

### Resting State Paradigm

Participants were presented with a white fixation cross on a gray background and asked to fixate while not doing any particular task. Resting state runs lasted 6 minutes each and for participants used in the resting state analysis, 14 had three runs and 4 had one run.

### MR Imaging

All scans were performed at Boston University’s Cognitive Neuroimaging Center on a Siemens 3-Tesla Prisma scanner with a 64-channel head coil. Structural images were collected for each participant with T1-weighted high-resolution magnetization-prepared rapid gradient multi-echo sequence (MP-RAGE, (van der Kouwe et al., 2008)) (1mm isotropic voxels, 176 slices, repetition time (TR) = 2530ms, echo times (TE) = 1.69, 3.55, 5.41 and 7.27ms, flip angle (FA) = 7°, field of view (FOV) = 256mm, GRAPPA (iPAT, [Griswold et al., 2002]) acceleration=3, TA=4min 23s). All blood-oxygen-level-dependent (BOLD) data were collected via T2*-weighted echo-planar imaging (EPI) pulse sequence that employed multiband RF pulses and simultaneous multi-slice (SMS) acquisition (Cauley et al., 2014; Feinberg et al., 2010; Moeller et al., 2010; Setsompop et al., 2012; Xu et al., 2013). Task fMRI (69 slices, TR=2000ms, TE=35ms, FA=80°, 2.2mm isotropic voxels, FOV=207mm, SMS=3, TA=7min 46s) and resting state fMRI (64 interleaved slices, TR=650ms, TE=34.8ms, FA=52°, 2.3mm isotropic voxels, FOV=207mm, SMS=8, TA=6min 9s) were also acquired. One participant’s resting state data was collected using the task fMRI sequence parameters, which was accounted for in the preprocessing pipeline. Pairs of reversed phase-encoded (anterior-posterior and posterior-anterior) spin echo field maps were also acquired for subsequent EPI de-warping.

### MRI Preprocessing

For ROI localization, structural and functional data preprocessing and general linear modeling were performed using Freesurfer (version 7.4.1, available at http://surfer.nmr.mgh.harvard.edu/) (Fischl, 2012). Anatomical reconstruction (recon-all) and registration (register-sess) were performed (Dale et al., 1999; Fischl et al., 2002) with human quality control inspection. Preprocessing included removal of non-brain tissue using a hybrid watershed/surface deformation procedure (Ségonne et al., 2004), automated Talairach transformation, intensity normalization (Sled et al., 1998), tessellation of the gray matter white matter boundary, automated topology correction (Fischl et al., 2001; Segonne et al., 2007), and surface deformation following intensity gradients to optimally place the gray/white and gray/cerebrospinal fluid borders at the location where the greatest shift in intensity defined the transition to the other tissue (Dale et al., 1999) (mkbrainmask-sess). Functional data preprocessing included motion correction (mc-sess), FSL’s “topup” and “applytopup” functions (Jenkinson et al., 2012; Smith et al., 2004) for field map distortion correction, slice-time correction (stc-sess), individual volume to fsaverage surface transformation (163,842 vertices) and smoothing with a 5mm full width at half max (FWHM) Gaussian kernel (rawfunc2surf-sess). General linear modeling included one regressor per stimulus condition convolved with the SPM canonical hemodynamic response function, as well as a nuisance regressor for linear drift removal, and 12 motion correction regressors per run (3 translation, 3 rotation, and their derivatives).

For functional connectivity analysis, anatomical and functional data were preprocessed using CONN (release 22.v2307) (Whitfield-Gabrieli & Nieto-Castanon, 2012) and SPM (release 12.7771) (Penny et al., 2011). The preprocessing pipeline consisted of slice timing correction, field map correction, outlier detection, direct coregistration to structural, resampling of functional data within the cortical surface, and smoothing of surface-level functional data. Slice-time correction was performed following SPM STC procedure (Sladky et al., 2011), using sinc temporal interpolation to resample each slice BOLD timeseries to a common mid-acquisition time. Outside of CONN, fieldmap distortion correction was applied using FSL’s “topup” and “applytopup” functions (Jenkinson et al., 2012; Smith et al., 2004). Back inside CONN, potential outlier scans were identified using ART (Whitfield-Gabrieli & Nieto-Castanon, 2011) as acquisitions with framewise displacement above 1 mm or global BOLD signal changes above 5 standard deviations (Power et al., 2014), and a reference BOLD image was computed for each subject by averaging all scans excluding outliers. Functional and anatomical data were coregistered using SPM intermodality coregistration procedure (Ashburner & Friston, 1997) with a normalized mutual information objective function (Studholme et al., 1998). Functional images were resampled at the cortical surface by averaging the functional data from ten locations between the pial and white matter cortical surfaces of each subject at each individual vertex in FreeSurfer fsaverage level-8 tessellation, with 163,842 vertices per hemisphere. Surface-level functional data were smoothed using 4 iterative diffusion steps, approximately a 3mm FWHM smoothing kernel within the cortical surface (Hagler et al., 2006). Last, anatomical data were segmented into grey matter, white matter, and CSF tissue classes using SPM unified segmentation and normalization algorithm (Ashburner, 2007; Ashburner & Friston, 2005) with the default IXI-549 tissue probability map template.

In addition, functional data were denoised using a standard CompCor (Behzadi et al., 2007) denoising pipeline including the regression of potential confounding effects characterized by white matter timeseries, CSF timeseries, motion parameters and their first order derivatives, outlier scans (Power et al., 2014), session and task effects and their first order derivatives, grey matter timeseries (Behzadi et al., 2007), and linear trends within each functional run, followed by bandpass frequency filtering of the BOLD timeseries between .008 Hz and .09 Hz. CompCor (Chai et al., 2012) noise components within white matter and CSF were estimated by computing the average BOLD signal as well as the largest principal components orthogonal to the BOLD average, motion parameters, and outlier scans within each subject’s eroded segmentation masks.

Task and resting state runs were rejected if there was more than 1mm average head displacement from midpoint position during the run (calculated using Freesurfer’s “mc-sess”). Twelve runs were rejected (7.6%), leading to one full subject rejection in the task dataset, and one full subject rejection in the resting state dataset. Thus, our task-based analyses contained twenty-one participants while our resting state analyses contained eighteen participants.

### Localizing Visual, Auditory, and Supramodal Regions of Interest in Individual Subjects

Visual-biased, auditory-biased, and supramodal WM ROIs in the present study were initially identified in prior studies with different data sets (Michalka et al., 2015; Noyce et al., 2022; Tobyne et al., in press). The present analysis follows the within-subject approach used in these earlier studies to define ROIs at the individual subject level. Prior work (Noyce et al., 2022; Tobyne et al., 2025) from our laboratory identified three visual-biased frontal lobe regions per hemisphere – superior precentral sulcus (sPCS), inferior precentral sulcus (iPCS), and mid-inferior frontal sulcus (midIFS) – and five auditory-biased frontal lobe regions per hemisphere – caudomedial superior frontal gyrus (cmSFG), transverse gyrus intersecting the precentral sulcus (tgPCS), caudal inferior frontal sulcus / gyrus (cIFS/G), central operculum (CO), and frontal operculum (FO). In occipitoparietal cortex we observed a large contiguous region that preferred visual processing (pVis), while in lateral temporal lobe we observed a large contiguous region that preferred auditory processing (pAud). Given the focus on frontal lobe regions, no attempt was made to isolate smaller, more specific function regions within pVis or pAud and the data contrasts were not conducive to subdivision of these contiguous regions. Additionally, recent work from the laboratory identified five supramodal frontal lobe regions in each hemisphere (Tobyne et al., 2025). Following a similar logic, we identify those five regions and one additional region in anterior cingulate cortex that emerged from the analysis. These supramodal regions are pre-supplementary motor area (preSMA-sm), anterior cingulate cortex (ACC-sm), anterior insula (aINS-sm), middle inferior frontal sulcus / middle frontal gyrus (midIFS/MFG-sm), superior precentral sulcus (sPCS-sm), and inferior precentral sulcus (iPCS-sm). Although five of these supramodal regions (not ACC-sm) lie in close proximity to visual-biased and/or auditory-biased regions, ROIs did not overlap in individual subjects. The full names, abbreviations, average MNI centroids, and average surface areas (mm^2^) for each ROI can be found in Table 1. All ROI creation was done after transforming the contrasts from individual cortical space to fsaverage. In total, 672 individual ROIs were identified for this study (16 ROIs x 2 hemispheres x 21 participants). The identification of such a large number of functional ROIs presents challenges, especially in frontal cortex where regions tend to be small and activation levels can be modest. Moreover, since the focus of this study is on functional connectivity measures, we required that each ROI contain a minimum of 100 cortical surface vertices. Our approach uses group-constrained ROIs to define subject-specific ROIs, paralleling an approach first used to define language-network regions (Fedorenko et al., 2010). Our ROI definition process is outlined in Figure 2, and the specifics are described below.

**Figure 2.**
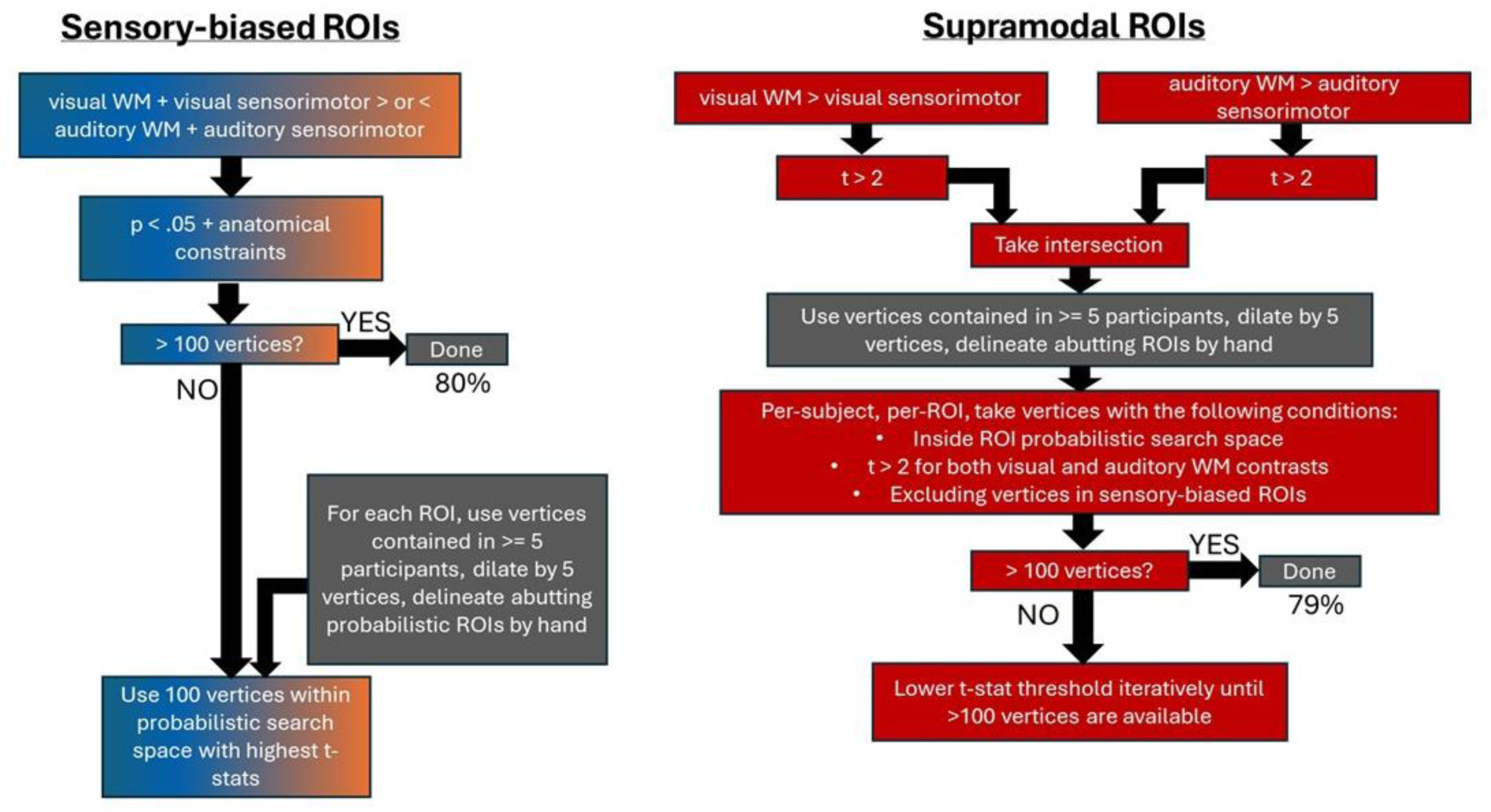
Flow chart summarizing the steps in defining the ROIs used in this study. 32 ROIs were defined for each individual participant. In cases where the subject-level data alone yielded a small ROI, probabilistic search spaces were used in conjunction with the subject-level data to yield ROI definitions of at least 100 vertices.

**Table 1.** Abbreviations, full names, sensory-bias, centroid MNI coordinates, and surface area for each ROI. The centroids were calculated using MNE’s “center_of_mass” function on each participant’s anatomical label in fsaverage space, then converted from vertex index to MNI coordinates using MNE’s vertex_to_MNI function. The coordinates were then averaged across participants. The surface area was calculated in individual subject cortical space using Freesurfer (mris_anatomical_stats).

| Abbreviation | Sensory Bias | ROI Name | MNI centroid (lh, rh) | Surface Area (mm<sup>2</sup>) (lh, rh) |
| --- | --- | --- | --- | --- |
| sPCS | Visual | Superior precentral sulcus | (-35.5, -7.4, 45.6),<br>(36.9, -5.7, 46.5) | 242±200, 328±281 |
| iPCS | Visual | Inferior precentral sulcus | (-45.6, 0.8, 32.1),<br>(43.2, 2.8, 29.8) | 319±265, 475±396 |
| midIFS | Visual | Middle inferior frontal sulcus | (-39.1, 29.3, 19.2),<br>(42.0, 29.7, 17.9) | 116±72, 182±196 |
| pVis | Visual | Posterior visual | (-29.2, -88.8, 3.0),<br>(30.8, -87.5, 3.6) | 8388±3055, 9493±3145 |
| tgPCS | Auditory | Transverse gyrus intersecting precentral sulcus | (-49.4, -6.7, 45.8),<br>(51.7, -3.8, 41.9) | 137±93, 142±79 |
| cIFS/G | Auditory | Caudal inferior frontal sulcus/gyrus | (-42.8, 13.7, 22.4),<br>(41.7, 15.1, 22.6) | 218±176, 178±122 |
| CO | Auditory | Central operculum | (-57.5, -5.1, 14.3),<br>(59.0, -3.7, 13.0) | 189±117, 170±146 |
| FO | Auditory | Frontal operculum | (-42.9, 30.8, 1.1),<br>(48.0, 29.5, 3.1) | 259±202, 209±152 |
| cmSFG | Auditory | Caudomedial superior frontal gyrus | (-7.0, 3.5, 61.3),<br>(7.9, 6.3, 62.1) | 102±96, 91±48 |
| pAud | Auditory | Posterior auditory | (-52.4, -24.0, 2.2),<br>(58.6, -18.2, 3.6) | 2979±708, 2864±646 |
| preSMA-sm | Supramodal | Supramodal pre-superior motor cortex | (-9.6, 14.9, 47.7),<br>(8.5, 22.0, 44.7) | 459±278, 568±321 |
| ACC-sm | Supramodal | Supramodal anterior cingulate cortex | (-12.5, 25.2, 30.1),<br>(12.6, 26.5, 26.4) | 89±44, 145±96 |
| aINS-sm | Supramodal | Supramodal anterior insula | (-28.3, 24.1, 3.5),<br>(29.9, 25.0, 3.7) | 526±300, 619±326 |
| sPCS-sm | Supramodal | Supramodal superior precentral sulcus | (-27.6, -4.4, 45.2),<br>(27.3, -2.4, 48.6) | 470±398, 471±446 |
| iPCS-sm | Supramodal | Supramodal inferior precentral sulcus | (-39.9, 2.5, 27.5),<br>(40.6, 9.8, 23.3) | 383±371, 571±509 |
| mid-IFS/MFG-sm | Supramodal | Supramodal mid-inferior frontal sulcus / middle frontal gyrus | (-42.2, 28.1, 25.9),<br>(42.1, 36.3, 23.3) | 344±278, 635±484 |

Visual and auditory frontal and posterior ROIs were defined at the individual subject level by directly contrasting visual WM and sensorimotor control blocks with auditory WM and sensorimotor control blocks. This contrast was liberally thresholded at p < .05, uncorrected in order to maximally capture vertices showing a bias for auditory or visual stimuli in individuals. Cortical significance maps were used in combination with anatomical expectations from previous studies (Michalka et al., 2015; Noyce et al., 2017, 2022; Tobyne et al., 2025) to define 5 bilateral frontal auditory-biased ROIs (tgPCS, cIFS/G, CO, FO, cmSFG), 3 bilateral frontal visual-biased ROIs (sPCS, iPCS, midIFS), and 2 large bilateral posterior sensory cortex ROIs (pVis, pAud). Each ROI was required to lie in or near the expected anatomical region (e.g., on the superior precentral sulcus), and exhibit the expected contrast (e.g., visual > auditory, p < .05).

Supramodal ROIs were defined for each individual by first directly contrasting visual WM with visual sensorimotor blocks, then separately contrasting auditory WM with auditory sensorimotor blocks. Then both contrasts were thresholded at t > 2 and their intersection was taken. A group-level probabilistic mask was then made from vertices contained in at least 5 individuals’ intersection masks. This probabilistic mask clearly segmented into 6 bilateral cortical clusters (Figure 3, red outline), similar to domain-general frontal regions localized in Assem et al. (2020) and Crittenden & Duncan (2014). We refer to these as supramodal (sm) ROIs (aINS-sm, ACC-sm, preSMA-sm, sPCS-sm, iPCS-sm, midIFS/MFG-sm) because they are active for both auditory and visual WM. The group-level probabilistic mask was then dilated by 5 vertices to create a search space for each supramodal ROI that better captures individual variability in location. Abutting supramodal ROI search spaces were delineated by hand at natural constriction points (Figure 3, red outline). Then, individualized ROIs were created by taking the individual auditory WM t >2, and visual WM t > 2 intersection vertices within each probabilistic supramodal ROI. For each subject, any vertices included in their sensory-biased ROIs were excluded from their supramodal ROIs.

**Figure 3.**
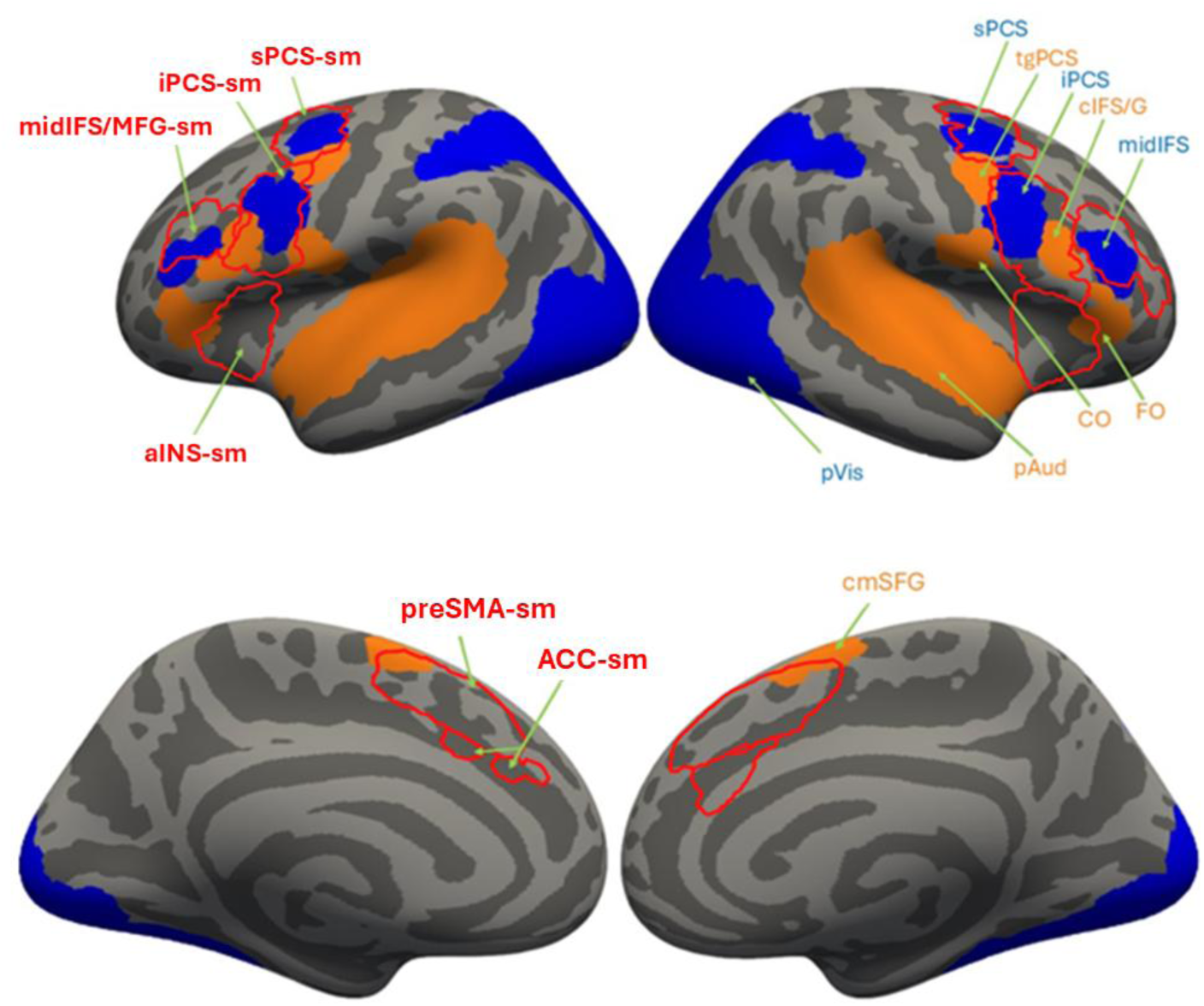
Lateral and medial left and right hemispheres with visual-biased probabilistic ROI search spaces in blue, auditory-biased probabilistic ROI search spaces in orange, and supramodal probabilistic ROI search spaces outlined in red. Name labels for the sensory-biased ROIs are shown only on the right hemisphere while supramodal ROI labels are shown only on the left hemisphere. Note that although there is overlap between the probabilistic supramodal and other probabilistic search spaces, the individual-subject ROIs never overlap.

Across both sensory-biased and supramodal, 21.5% of ROIs (138/672) identified had fewer than 100 vertices. We replaced these ROIs with the strongest 100 vertices within the probabilistic search space for the ROI. To expand the sensory-biased ROIs with fewer than 100 vertices, we first created probabilistic sensory-biased ROIs (Figure 3, blue and orange) with the same process used to create the supramodal probabilistic ROIs. Then we used the 100 vertices with the largest t-stats within the probabilistic ROI to replace the sensory-biased ROIs with fewer than 100 vertices. Only 7 sensory-biased ROIs (∼1%) contained any vertices with t-stats in the opposite direction than expected. For the supramodal ROIs with fewer than 100 vertices, the auditory WM > auditory sensorimotor and visual WM > visual sensorimotor t-stat thresholds were incrementally lowered together until 100 vertices were included, excluding any vertices in the sensory-biased ROIs. Across all subjects, only 4 supramodal ROIs (<1%) contained any vertices with negative t-stats.

Two participants were rejected from all analyses due to inability to draw more than 75% of ROIs at a liberal p < .05 uncorrected threshold; for one there was low overall activation (ex. no significant activity in the auditory cortex), while for the other, activity was heavily skewed towards visual and sensorimotor control conditions. This left 21 participants (18 resting state).

### Analysis Overview

We employed several individual-subject functional connectivity analyses to investigate the resting state and task-based network organization of sensory-biased and frontal supramodal ROIs. To establish robust network organization at rest (N=18) we used hierarchical clustering analysis (HCA), then used linear mixed effects models (LME) on ROI-to-ROI correlations grouped by network to establish connectivity patterns between the networks. To reveal which specific ROIs from each network were driving connectivity, we used threshold-free cluster enhancement (TFCE) on the ROI-to-ROI correlations. To then understand how network-network and ROI-ROI connectivity differentially change with visual and auditory WM task demands (N=21), we used generalized psychophysiological interaction (gPPI) analysis. The PPI coefficients were grouped by network and used in LME analysis to see how network connectivity might restructure during visual and auditory WM tasks. Again, to reveal which specific ROIs from each network were driving these changes, we used TFCE on the ROI-to-ROI PPI coefficients. The resting state and task-based network-based LME connectivity analyses were the primary hypothesis-driven analyses while the ROI-specific TFCE analyses were post-hoc.

### Resting State Functional Connectivity Analysis

HCA was used to determine which ROIs had the most correlated activity. CONN Toolbox computed complete-linkage clustering (Sørensen, 1948) with optimal leaf ordering (Bar-Joseph et al., 2001) based on functional similarity only (contribution of anatomical proximity was set to zero) (Nieto-Castanon, 2020). The number of clusters was selected by maximizing the difference between normalized number of clusters and the normalized inverse distance metric (i.e. analogous to choosing the “elbow” of the curve plotting the number of clusters against the distance metric). These clusters were considered as functional networks in the following analyses.

To assess resting-state connectivity between networks found via HCA, we first used CONN Toolbox to compute ROI-to-ROI connectivity using Fisher-transformed bivariate correlation coefficients from a general linear model (weighted-GLM) for each pair of ROIs separately, thus characterizing the association between their BOLD signal timeseries. To compensate for potential transient magnetization effects at the beginning of each run, individual scans were weighted by a step function convolved with the SPM canonical hemodynamic response function and rectified (Nieto-Castanon, 2020). We then used these ROI-to-ROI correlation coefficients as the dependent variable in an LME (r), with network connection type as the fixed effect (n), and subject (s) and hemisphere (h) as full random effects (intercept, slope, and covariance). The model specification in Wilkinson notation formatted for Matlab is as follows:

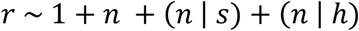

Network connection type was assigned based on which HCA-determined networks the ROIs were in; for example, the correlation coefficient for sPCS and sm-aINS would be labeled as a “*frontal visual*←→*supramodal*” network connection. Because this analysis is GLM-based, there were separate coefficients for reciprocal connections, which were averaged to represent the directionally agnostic connectivity between two ROIs. Next, estimated marginal means (EMMs) and standard errors of connectivity for each connection type were computed from the LME coefficients and used in Wald-tests for significance (Hartman, 2019/2023). Estimated effect sizes for coefficients were calculated by taking the difference between EMMs (or the difference between a single EMM and the reference point of zero) and dividing by the standard deviation of the sample response variable (r) (Hedges, 2007). P-values were FDR-corrected for significance at p < .05. The robustness of this method relies on using fully specified random effects (intercept, slope, and covariance) of subject and hemisphere to account for hemispheric- and subject-level effects and enhance generalizability (Barr et al., 2013), and FDR-correction for multiple comparisons.

Assessing significance of individual ROI-to-ROI connectivity is complicated by the multiple comparisons problem due to the large number of ROIs (496 comparisons among 32 ROIs), thus we used the nonparametric spatial clustering method TFCE to reduce the number of comparisons being made. This method has the additional advantage of increased sensitivity by avoiding use of an arbitrary cluster-forming threshold, and it easily lends itself to permutation testing for significance (Smith & Nichols, 2009). We used CONN Toolbox to first estimate a separate GLM for each connection, creating a group-level ROI-to-ROI connectivity matrix, then used the CONN Toolbox implementation of TFCE to determine clusters of significant connectivity among ROIs, using HCA determined network membership as the adjacency matrix. Thus, connections (edges) containing ROIs in the same network are considered adjacent for TFCE clustering without regard for the spatial distance between them (this would introduce spatial proximity bias). Significant clusters of connections were determined using permutation testing of 10,000 residual-randomization iterations with FDR correction of TFCE-scores at p < .05 (Nieto-Castanon, 2020). The robustness of this method relies on avoiding the use of a hard cluster-level threshold, nonparametric permutation testing (Eklund et al., 2016, 2019), and FDR-correction for multiple comparisons.

### Task-Based Functional Connectivity Analysis

To assess the changes in connectivity during visual and auditory WM tasks, we used gPPI analysis implemented in CONN Toolbox, using individual-specific ROIs. Although in group analyses, task modulations account for a small portion of the variance in functional connectivity (e.g., Cole et al., 2021; Gratton et al., 2018; Krienen et al., 2014; Smith et al., 2009), the application of individual-specific methods yield substantially greater power (Gratton et al., 2018). PPI assesses changes in connectivity between 2 ROIs due to particular experimental conditions using bivariate regression. Each regression model estimates the variance in one ROI BOLD timeseries explained by 1) terms for each task effect convolved with a hemodynamic response, 2) the seed ROI BOLD timeseries, and 3) the interaction term, which is the product of terms 1 and 2. Thus, by estimating the coefficients for term 3 (the interaction term or PPI), we can quantify the amount of variance in one ROI signal explained by the interaction between another ROI and each task. Though the ROIs were defined from the task data set, the PPI analysis is non-circular because 1) task effects and their derivatives are regressed out in the denoising step, and 2) PPI analysis assesses the predictability of one ROI’s signal by another ROI’s signal under a specific condition *over and above* the variability explained by the seed ROI’s signal alone (term 2) and the task effect alone (term 1); that is, the PPI analysis only detects effects that are orthogonal to the main effect of the task (Friston et al., 1997; McLaren et al., 2012; O’Reilly et al., 2012). Additionally, gPPI is shown to estimate context-modulated functional connectivity especially well under block-designs compared to beta-series methods and sPPI (Cisler et al., 2014).

The gPPI analysis was run for every ROI-to-ROI connection and PPI coefficients were used to form 2 PPI matrices, one for the change in connectivity between ROIs during visual WM and one for auditory WM. Note that PPI coefficients represent the impact of a specific experimental condition on connectivity between ROIs compared to a “baseline” of all other conditions (which here included fixation and sensorimotor blocks). Next, these matrices were subtracted in order to get the difference in connectivity between visual and auditory WM. This matrix was then evaluated in the same way as the connectivity coefficient matrix in the above LME and TFCE analyses. Thus, the LME analysis with the PPI data showed how the network-level connectivity differed between visual and auditory WM tasks, while the TFCE analysis showed how ROI-level connectivity differed between the visual and auditory WM tasks.

### Assessing Overlap Between Supramodal and Domain-General Frontal Regions

In order to assess the similarity between our frontal supramodal ROIs and the domain-general frontal ROIs from the literature (Assem et al., 2022), we used the Sørensen-Dice similarity coefficient (Dice, 1945; Sørensen, 1948) which simply takes 2 times the number of vertices in common between the two ROIs and divides by the sum of the number of vertices in both ROIs, thus quantifying the spatial similarity between the ROIs:

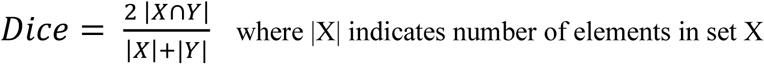

We assessed only the ROI pairs for which the probabilistic supramodal search space visually intersected with the domain-general ROIs (derived from the HCP-MMP1.0 parcellation) when projecting both on the fsaverage cortex (Supplementary Figure 1). Subsequently, we used the individually defined supramodal ROIs to assess overlap with the fsaverage-projected domain-general ROIs for each participant.

### Predicting Working Memory Precision from Dynamic Network Connectivity

After the primary analyses, to evaluate the relationship between subject-level behavioral task precision and task-related changes in connectivity among networks during each task, we fit linear regression models using task precision as the dependent variable and a subset of network-level PPI difference terms as the independent variables. Working memory precision for each participant was defined by the average staircase threshold obtained. Separate staircasing was done for visual and auditory stimuli. The staircasing procedures sought to equate task difficulty (accuracy) across modalities and across individuals by adjusting the size of the frequency difference between a non-match stimulus and a match stimulus. Staircase thresholds were recorded as Weber fractions or percentage changes from the baseline. Finer thresholds (i.e., smaller frequency difference between stimuli) indicate finer WM precision. Thus, if a participant performed well (poorly) on the task, their difficulty staircase would increase (decrease), and it would be harder (easier) to tell the difference between the first and second stimulus in a non-match trial. To predict task precision in the *visual* WM task, we averaged the PPI coefficient differences across hemispheres and across network connections with significant increases in connectivity during visual WM compared to auditory WM. Similarly, to predict task precision in the *auditory* WM task, we averaged the PPI coefficient differences across hemispheres and network connections with significant increases in connectivity during auditory WM compared to visual WM. Our reasoning was that the network-network connections which showed significant changes during the task were most likely to impact behavioral performance.

## Results

Subjects (N=21) performed 2-back visual working memory and 2-back auditory working memory tasks during fMRI (see Figure 1; Methods). Task accuracy did not differ significantly between visual and auditory WM tasks (visual WM=68.1% auditory WM=64.8%, t(20)=1.91, p=.08) (Supplementary Figure 2).

We examined the 2-back WM data to define 32 ROIs (16 ROIs per hemisphere) for each individual subject (See Figures 2; Methods). Prior work from our laboratory identified three visual-biased, five auditory-biased, and five supramodal regions in frontal lobe in both hemispheres (Noyce et al., 2022; Tobyne et al., 2025). Individual subject ROIs were compiled to create probabilistic ROI maps, which are displayed in Figure 3. Individual subject ROIs are presented in Figure 4 and in Supplementary Figure 3. ROI names, MNI coordinates and region sizes are summarized in Table 1.The mean MNI coordinates aligned well with those regions defined in prior studies (Noyce et al., 2022; Tobyne et al., 2025), even though the subjects and stimuli differed across datasets (Supplementary Tables 1, 2). Supramodal regions, which were defined here based on the two-way intersection of visual WM and auditory WM activation, tended to be larger than those defined by a three-way intersection (visual, auditory, and tactile WM) in the prior analysis (Tobyne et al., 2025), but otherwise appear well-aligned.

**Figure 4.**
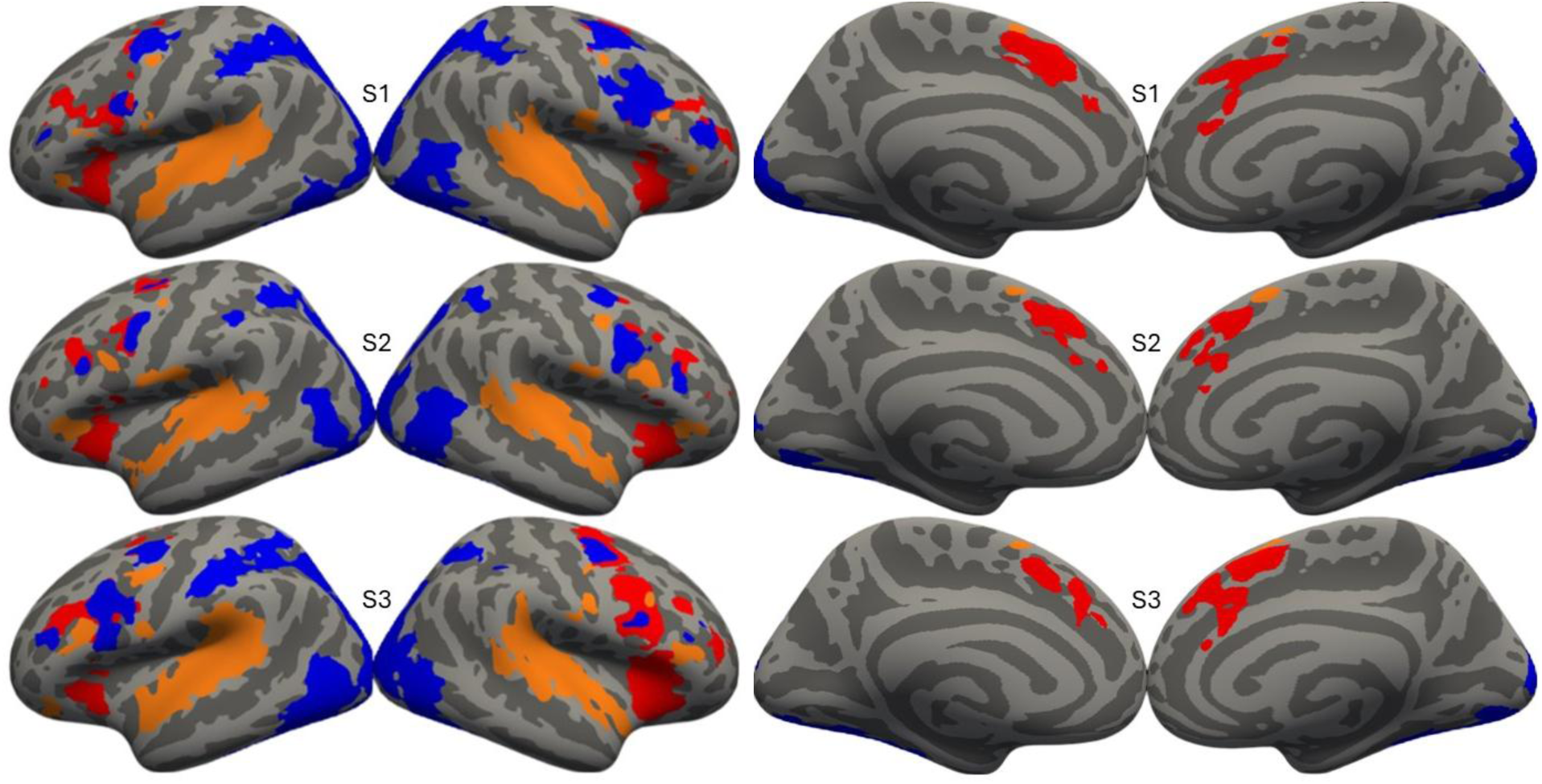
Three individual participants’ sensory biased and supramodal ROIs on the fsaverage lateral and medial cortical surfaces. Blue ROIs indicate visual bias, orange indicates auditory bias, and red indicates supramodal ROIs. ROIs for all subjects can be found in Supplementary Figure 3.

We note that the supramodal ROIs we functionally defined bear a striking anatomical similarity to a large subset of the domain-general network identified in broad task batteries (Assem et al., 2020; Fedorenko et al., 2012). To examine the anatomical relationship between the present set of supramodal WM ROIs and the previously defined domain-general ROIs, we performed Dice coefficient and percent overlap analyses between our individual-subject supramodal ROIs and the MMP 1.0 multi-modal parcels (Glasser et al., 2016) that are identified as the domain-general network (Assem et al., 2020). Only pairs of ROIs in which the supramodal ROI search space overlapped with a domain-general ROI were assessed. Five of the six supramodal ROIs identified here heavily overlapped (>50%) with the frontal domain-general ROIs (see Supplementary Figure 1; Supplementary Table 3). This gives evidence that our supramodal WM ROIs may constitute a substantial subset of domain-general cortex.

### Resting-State Functional Connectivity: Auditory, Visual, and Supramodal Networks Form Distinct but Interconnected Resting State Networks

We examined network organization amongst 32 ROIs using resting-state fMRI data to establish baseline, task-free connectivity. Hierarchical Clustering Analysis (HCA) of the seed-to-seed functional connectivity matrix revealed 4 intra-connected groups of ROIs (see Methods). One network consisted of bilateral visual-biased frontal ROIs (sPCS, iPCS), another consisted of the bilateral auditory-biased frontal and posterior ROIs (tgPCS, cIFS/G, CO, FO, cmSFG, pAud), and another consisted of all the bilateral supramodal frontal ROIs plus a single visual-biased frontal ROI (aINS-sm, preSMA-sm, ACC-sm, sPCS-sm, iPCS-sm, midIFS/MFG-sm, midIFS). The final cluster consisted of the single visual-biased posterior bilateral ROI, posterior visual (pVis). We refer to these networks as *frontal visual*, *frontal+posterior auditory*, *supramodal*, and *posterior visual* networks respectively (see Supplementary Figure 4 for full ROI connectivity matrix). The HCA results largely agreed with previous results that showed visual-biased and auditory-biased frontal ROIs formed separate networks (Michalka et al., 2015; Tobyne et al., 2017; Noyce et al., 2022) and extended previous results by showing that supramodal ROIs formed another separate network, with the exceptions that pAud clustered with the frontal auditory ROIs and midIFS clustered with the *supramodal* network instead of the *frontal visual* network.

To characterize the connectivity between networks, we used linear mixed effects (LME) analysis of correlations between ROIs in each network (see Methods). These connectivity patterns and the mean correlations are shown in Figure 5a (also see Supplementary Note 2: LME Assumption Checks). In general, we saw two distinct streams of positive connectivity, a ‘visual stream’ between the *posterior visual*, *frontal visual*, and *supramodal* networks, and separately, an ‘auditory stream’ between the *posterior+frontal auditory* and *supramodal* networks. Effect sizes for significant connectivity strength between networks ranged from small to large with a mean Cohen’s d of .71 (SEs, p-values, and effect sizes are shown in Supplementary Table 4). The connectivity between *frontal visual* ←→ *supramodal* (M=.15, SE=.02) was significantly greater (W(1)=4.1, p<.001, Cohen’s d=.33) than between the *frontal+posterior auditory* ←→ *supramodal* networks (M=.06, SE=.02). This finding replicated and extended the sensory network asymmetry previously reported (Tobyne et al., 2017; Noyce et al., 2022) by demonstrating that the visual stream exhibited greater functional connectivity with supramodal regions than did the auditory stream. Note that this functional connectivity asymmetry existed even though the supramodal ROIs did not overlap with sensory-biased ROIs. It is also of note that there was only a single point of significant positive connectivity between the visual and auditory networks, and that connection between *frontal visual* ←→ *frontal+posterior auditory* was weak (M=.04, SE=.02), indicating largely separate visual and auditory streams. Given that the nodes of the frontal visual and auditory networks are anatomically interleaved with each other and abut in many instances, the relative segregation of the modality-biased networks is remarkable.

**Figure 5.**
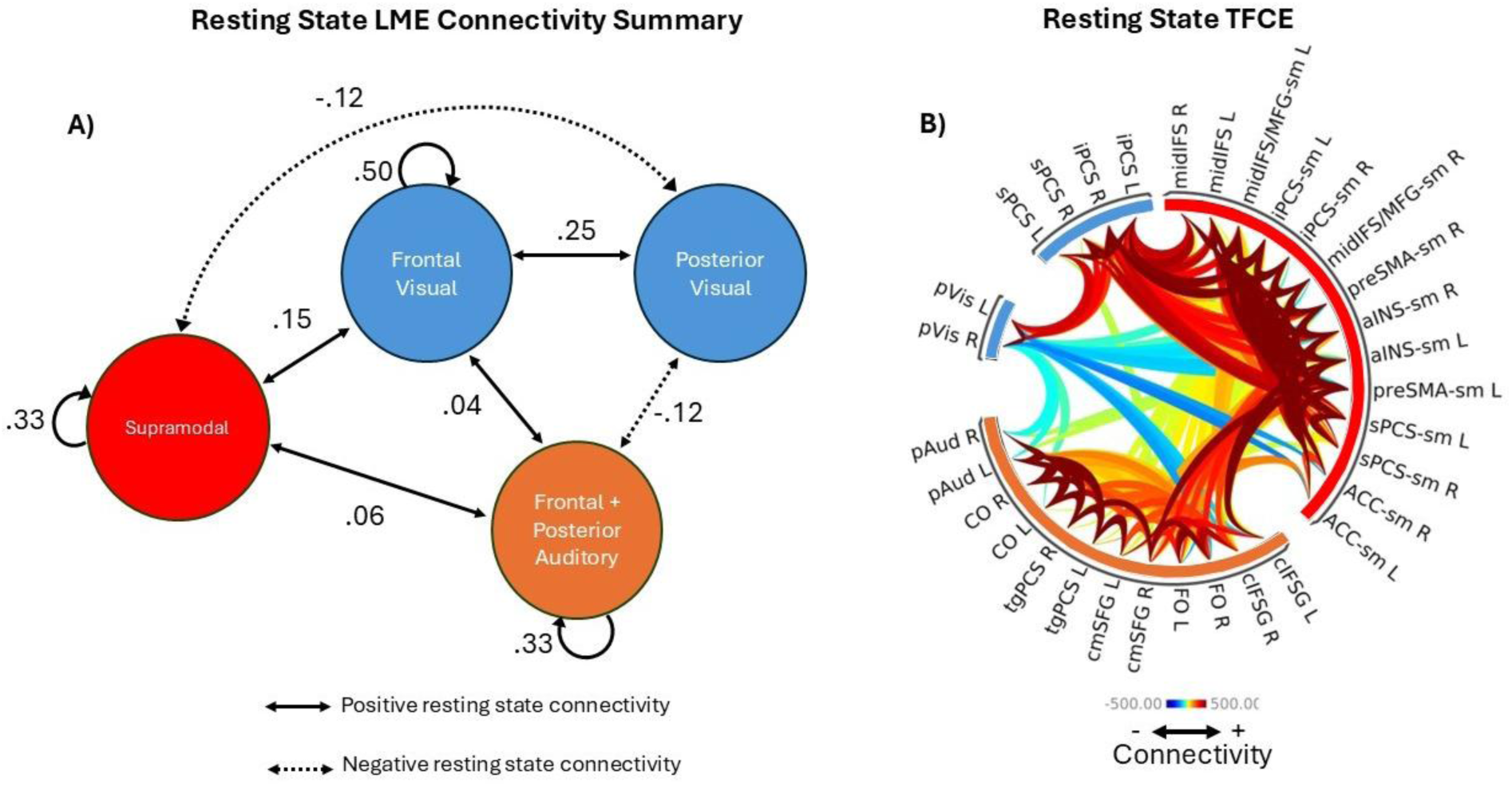
Resting-state connectivity results. A) Diagram summarizing the LME-derived significant resting state connectivity between ROI networks. The values shown next to each connection represent the mean correlation coefficient between ROIs in each network. Self-connection in the *posterior visual* network was not assessed because it only contained 1 (bilateral) ROI. B) Connectogram summarizing the TFCE-derived significant resting state connectivity between individual ROIs. Warm colors represent positive connectivity while cool colors represent negative connectivity. ROIs are grouped around the circle by HCA-derived networks.

We performed three sets of control analyses on the resting-state networks: 1) we repeated the LME analysis using only the ∼80% of ROIs defined robustly from individual subject data alone (Supplementary Table 4); 2) we repeated the LME analysis using a network grouping in which the *frontal+posterior auditory* network was replaced by separate *frontal auditory* and *posterior auditory* networks (See Supplementary Table 5); and 3) we included ROI-to-ROI distance as a random-effect in our LME analysis, to address concerns about distance-dependent correlations. Critically, in all 3 analyses the connectivity between supramodal and frontal visual networks was significantly greater (p<.001) than between supramodal and frontal auditory networks. Full details are described in Supplementary Note 1: Resting-State Network Controls.

To assess connectivity between specific ROIs within each network, we next used TFCE of the seed-to-seed functional connectivity with ROIs sorted using the HCA clustering results (see Methods). Results from the TFCE analysis are shown in Figure 5b where ROIs are grouped into their HCA-identified networks around the outside of the connectogram. Most clusters recapitulated the connectivity patterns observed in the LME analysis, but now with ROI-level specificity. For example, the negative connectivity between *posterior visual* ←→ *frontal+posterior auditory* networks was driven by negative connectivity of pVis with pAud, CO (L), and FO. We also saw that there was especially strong connectivity between ROIs in the *frontal visual* ←→ *supramodal* networks while connectivity between ROIs in the *auditory frontal+posterior* ←→ *supramodal* networks was present, but generally weaker.

### Task-Based Functional Connectivity: Auditory Working Memory Drives More Widespread Changes in Connectivity than Visual Working Memory

The above results demonstrated that: 1) largely distinct visual-biased and auditory-biased networks exist in the cortex; 2) each modality-biased network connected within the frontal lobe to a supramodal network; and 3) in the resting state, supramodal regions exhibited stronger functional connectivity with the visual network than with the auditory network. How can we reconcile the network asymmetry *at rest* with the apparent unbiased supramodal activation *during WM task performance*? We hypothesized that during auditory WM task performance the auditory stream increases its functional connectivity (compared to baseline) and does so to a greater extent than occurs for the visual stream during visual WM.

We first performed a network-level LME analysis of PPI coefficients (see Methods) and observed an increase in connectivity between the *frontal+posterior auditory* ←→ *supramodal*, *frontal+posterior auditory network* ←→ itself, and *frontal+posterior auditory* ←→ *frontal visual* networks during auditory WM, as compared to visual WM. We also saw an increase in connectivity between the *posterior visual* ←→ *frontal visual*, and *posterior visual* ←→ *supramodal* networks during visual WM, as compared to auditory WM (Figure 6a). Notably, there was no significant change in connectivity between the *frontal visual* ←→ *supramodal* networks during the two working memory conditions. It is also important to note that the increases in connectivity during auditory WM compared to visual WM were largely driven by an increase in connectivity during auditory WM (compared to baseline), whereas the visual>auditory WM increases in connectivity were driven largely by decreases in connectivity during auditory WM (compared to baseline) (Table 2). Estimated marginal means, SEs, p-values, and effect sizes for the change in connectivity between each network pair are shown in Supplementary Table 6. Effect sizes of significant differences ranged from small to medium with a mean Cohen’s d of .30. In order to ensure that the results were not unexpectedly biased by the inclusion of individual ROIs defined with guidance from the probabilistic maps, we repeated the above LME analysis in full using only the ∼80% of ROIs defined robustly from individual subject data alone. All significant and non-significant results remained that way (Supplementary Table 6). Additionally, to ensure the inclusion of the fixation condition in the baseline for gPPI did not bias the interpretation of WM modulation of connectivity, we repeated the gPPI LME analysis without the fixation condition in the baseline conditions and found that all significant and non-significant results remained that way (Supplementary Table 7).

**Figure 6.**
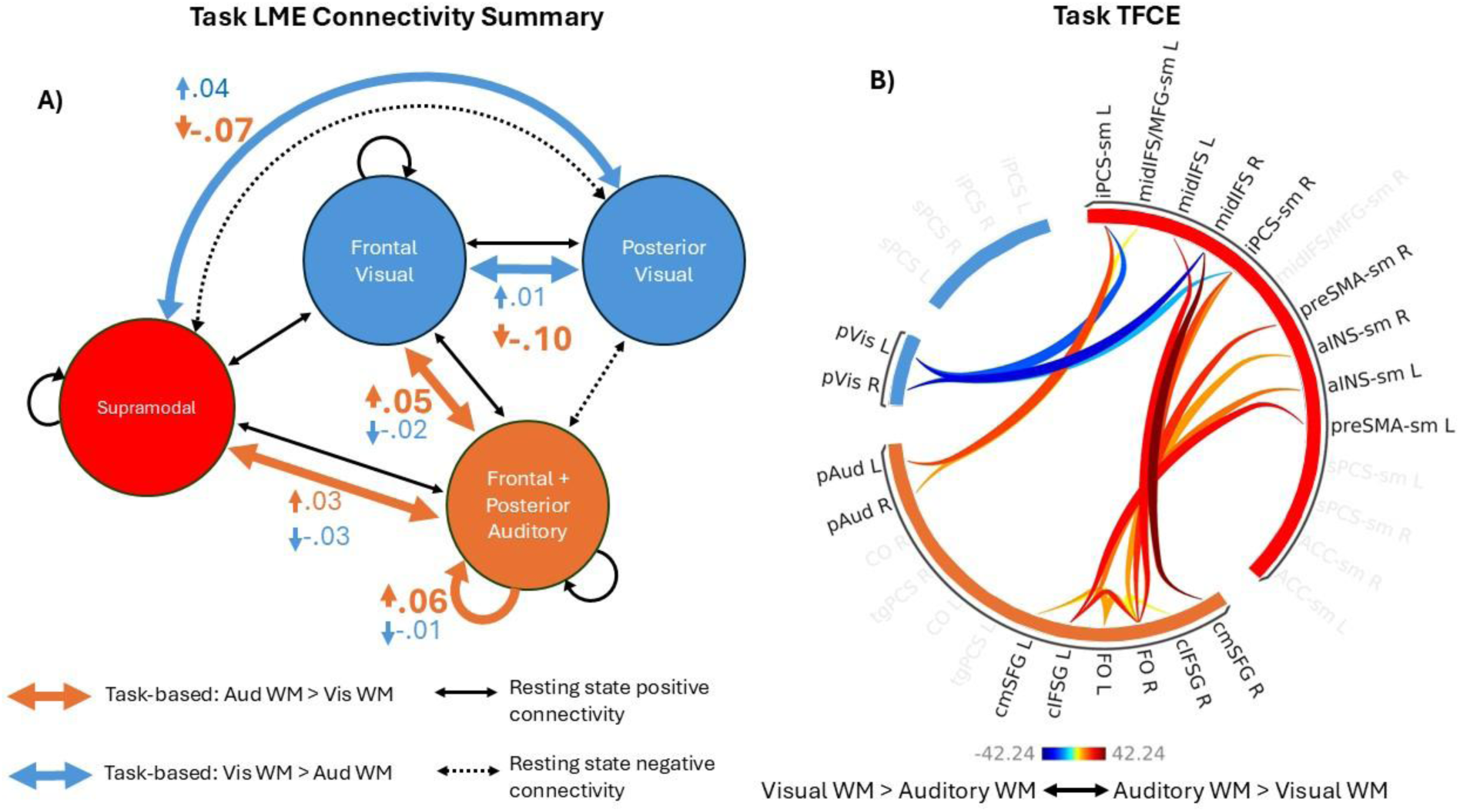
Task-based functional connectivity results. A) Diagram summarizing the LME-derived significant changes in connectivity between ROI networks during visual and auditory WM. The black arrows show significant resting-state connectivity from Figure 5, while the large colored arrows indicate change in connectivity. The values and small colored arrows shown next to the large colored arrows indicate the mean increase or decrease in PPI coefficients for the visual and auditory WM conditions between ROIs in each network. For example, the connectivity between *posterior visual* ←→ *frontal visual* networks increased under visual WM (large blue arrow) and was driven by a small (.1) increase in the visual WM condition and a large decrease (.10) in the auditory condition (small blue and orange arrows and values). Changes in self-connection in the *posterior visual* network were not assessed because the network only contained 1 (bilateral) ROI. B) Connectogram summarizing the TFCE-derived significant changes in connectivity between individual ROIs during visual and auditory WM. Warm colors represent auditory>visual changes in connectivity while cool colors represent visual>auditory changes in connectivity. ROIs are grouped around the circle by HCA-derived networks.

**Table 2.** Means±SEs, and p-values for the visual and auditory WM PPI terms for each network connection. Bold p-values indicate significance under FDR correction.

| Connection Type | Mean Visual WM PPI Beta | Mean Auditory WM PPI Beta | p-value |
| --- | --- | --- | --- |
| *post vis<->supramodal* | .041 ± .013 | -.069 ± .014 | **.001** |
| *post vis<->front vis* | .011 ± .021 | -.096 ± .024 | **.008** |
| *post+front aud<->post+front aud* | -.010 ± .007 | .061 ± .008 | **.001** |
| *post+front aud<->front vis* | -.015 ± .009 | .047 ± .009 | **.001** |
| *post+front aud<->supramodal* | -.027 ± .005 | .026 ± .005 | **.005** |
| *supramodal<->front vis* | -.014 ± .007 | .024 ± .008 | .123 |
| *front vis<->front vis* | -.047 ± .021 | -.015 ± .023 | .624 |
| *post+front aud<->post vis* | -.022 ± .014 | -.007 ± .015 | .583 |
| *supramodal<->supramodal'* | -.002 ± .006 | .012 ± .006 | .504 |

These findings suggest that auditory WM task performance caused more profound restructuring of connectivity between networks than did visual WM task performance. A post-hoc summary analysis of the changes in connectivity between networks in the proposed “visual stream” during visual WM was compared against the changes in connectivity between networks in the proposed “auditory stream” during auditory WM. Thus, a one-way ANOVA with random effects for subject and hemisphere was run comparing the average visual WM PPI terms for the following connections: *posterior visual* ←→ *frontal visual, frontal visual* ←→ *frontal visual, frontal visual* ←→ *supramodal*, and the average auditory PPI terms for the following connections: *frontal+posterior auditory* ←→ *frontal+posterior auditory, frontal+posterior* ←→ *supramodal*. There was a statistically significant difference showing larger average changes in connectivity across the “auditory stream” during auditory WM compared to baseline than average changes in connectivity across the “visual stream” during visual WM compared to baseline (F(1,125)=6.24, p=.014). The effect size (η^2^) was .04 indicating between a small and medium effect size.

We also assessed ROI-level changes in connectivity using TFCE on the seed-to-seed PPI coefficients with ROIs sorted using the resting state HCA clustering results (Figure 6b; see Methods). Again, we saw that the significant clusters of connectivity changes at the ROI-level matched well with the patterns of connectivity changes in the network-level analysis, but we gained specificity as to which ROIs were driving the changes. The majority of significant differences in connectivity between auditory and visual WM in ROI-ROI connectivity identified were the result of increased connectivity under auditory WM compared to baseline, again consistent with the idea that auditory WM leads to greater reorganization of functional connectivity within and between sensory-biased and supramodal networks than does visual WM.

Additionally, to see if the overall results were idiosyncratic to small changes in network clustering, we repeated the above LME analysis using *a priori* symmetric network grouping in which the midIFS is included in the *frontal visual* network and pAud is pulled out of the *frontal+auditory* network to form a *posterior auditory* network. The resulting changes in connectivity structure included an increase in connectivity between *posterior auditory* ←→ *frontal auditory* networks during auditory WM as compared to visual WM (Supplementary Table 5). Additionally, when separated, neither the *frontal auditory* network nor *posterior auditory* network reach significance in their increase in connectivity to the *supramodal* network during auditory WM compared to visual WM, although when combined in the previous analysis, they do, potentially indicating insufficient power to detect an effect when looked at individually. To investigate this further, we repeated the above TFCE analysis with the *a priori* symmetric network groupings and found that 20 of 24 (83.3%) significant ROI-ROI pairings from the original analysis remained in this analysis, with 6 ROI-ROI pairings between the *frontal auditory* ←→ *supramodal* networks increasing connectivity during auditory WM compared to visual WM, while zero ROI-ROI pairing between the *frontal visual* ←→ *supramodal* networks increased connectivity during visual WM compared to auditory WM (Supplementary Figure 5). Lastly, we reran the summary ANOVA analysis and still found a statistically significant difference showing larger average changes in connectivity across the “auditory stream” during auditory WM compared to baseline than average changes in connectivity across the “visual stream” during visual WM compared to baseline (F(1,125)=5.11, p=.026) with an effect size of η^2^=.03. This suggests that although splitting the *frontal* and *posterior auditory* networks dilutes detection power, the coarse level contrast of dynamic effects in the auditory and visual networks persists.

### Auditory WM Precision is Correlated with Changes in Connectivity Between Key Networks

In order to assess the behavioral impact of shifting connectivity between networks during auditory and visual WM tasks, we used linear regression with WM task precision as the dependent variable and the difference between visual and auditory PPI coefficients as the independent variables (see Methods). After the above PPI analyses were completed, we hypothesized that at the individual level, increases in connectivity between key networks during auditory WM would predict greater auditory WM precision. Those key network connections were selected based on which networks had significantly increased connectivity during auditory WM compared to visual WM in the previous group level analysis (*frontal+posterior auditory* ←→ *supramodal, frontal+posterior auditory* ←→ *frontal visual,* and *frontal+posterior auditory* ←→ *frontal+posterior auditory*). In line with this hypothesis, we found that changes in connectivity in these key network-level connections during auditory WM compared to visual WM significantly predicted behavioral precision on the auditory WM task (R^2^=.20, F(1,19)=4.81, β=-37.76, p=.0204; Figure 7).

**Figure 7.**
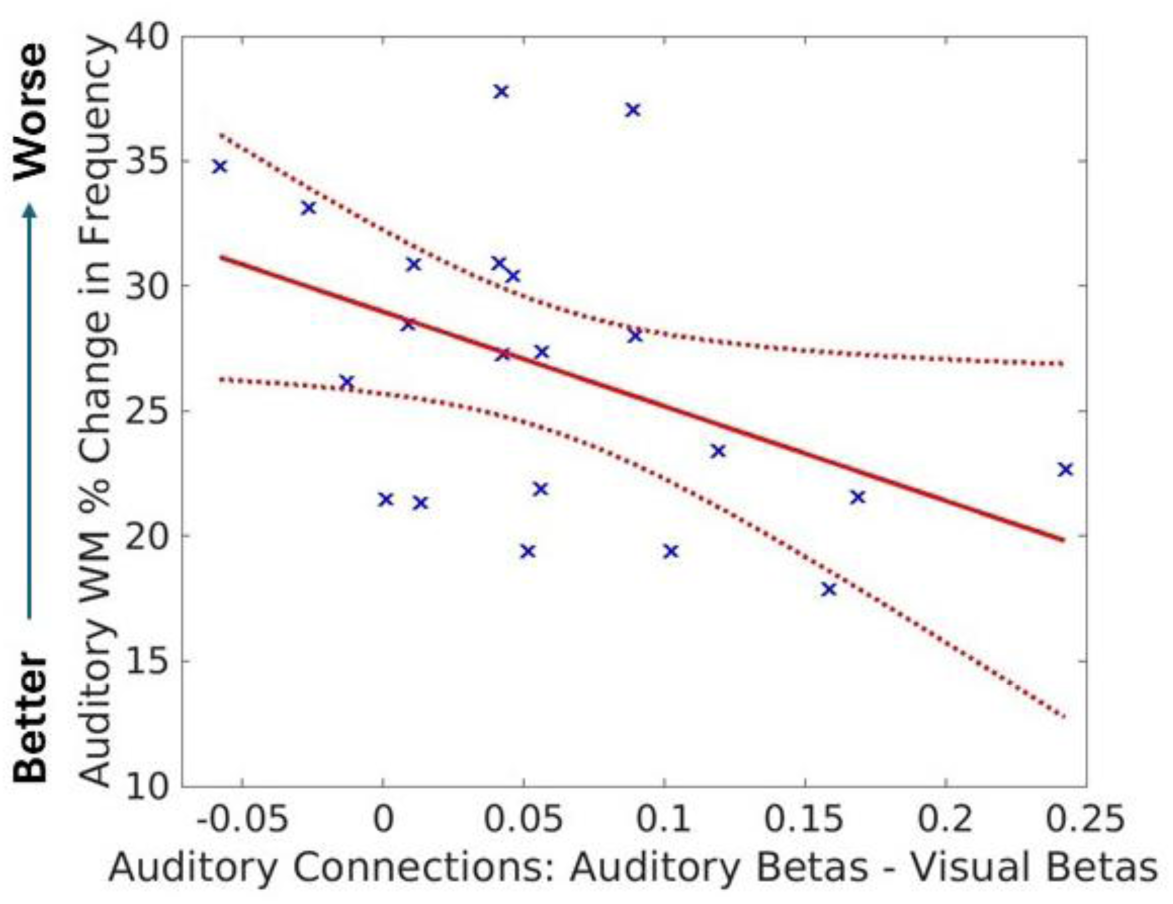
Auditory WM precision in individual participants correlated with the strength of the auditory network reconfiguration during the WM task. Each blue X represents a single participant’s data (N=21). The red line is the least squares line of best fit, and the dotted lines are 95% confidence bounds. The Y axis represents percent change in frequency between stimuli (lower indicates higher behavioral precision) Note that the x-axis represents the average difference in PPI betas for the 3 network-network connections with significantly increased connectivity during auditory WM compared to visual WM.

We also hypothesized that increases in connectivity between key networks (*posterior visual* ←→ *frontal visual* and *posterior visual* ←→ *supramodal*) during visual WM compared to auditory WM would *not* predict better visual WM task precision. This hypothesis was derived from the fact that visual WM elicited much smaller changes in connectivity between networks than auditory WM, possibly because frontal visual ROIs are already strongly connected to supramodal ROIs at rest. Mirroring the analysis done on the auditory WM task, this analysis examined the networks that had significantly increased connectivity during visual WM compared to auditory WM. In line with this hypothesis, we found that changes in connectivity in key network-level connections during visual WM compared to auditory WM did not predict behavioral precision on the visual WM task (R^2^=.01, F(1,19)=.25, β=-6.52, p=.626).

## Discussion

In this study we sought to understand the functional connectivity relationships among visual-biased, auditory-biased, and supramodal cortical ROIs, and how they change during visual and auditory WM tasks. Our results demonstrated that visual-biased, auditory-biased, and supramodal ROIs form largely discrete but interconnected resting state networks. We saw little positive connectivity between visual and auditory network streams indicating that there are relatively separated processing streams for visual and auditory sensory and WM information, even in the frontal cortex – with both streams potentially feeding into and/or receiving information from a modality-agnostic frontal supramodal network. This segregation is remarkable given that the nodes of these two networks are anatomically interleaved in frontal cortex. Several of our key findings focus on asymmetries between the visual-biased and auditory-biased networks. Resting-state analysis demonstrated that the *supramodal* network was more highly connected with the *frontal visual* network than with the *frontal+posterior auditory* network. This result held under several control analyses. This suggests that the *frontal visual* network has “privileged” communication with the *supramodal* network at rest. This idea is supported by the lack of detectable significant changes in functional connectivity between the *frontal visual* and *supramodal* networks during the visual WM task; perhaps the strong connectivity observed at rest is sufficient to support strong visual WM performance. In contrast, there was much greater reorganization of connectivity during the auditory WM task, including the recruitment of the *visual frontal* and *supramodal* networks, which also correlated with increased auditory WM precision in individual participants in a post-hoc analysis. No such changes were observed for the visual WM task. That is, although the auditory-biased network appeared disadvantaged at rest in its connectivity to the supramodal WM network, it was capable of greater dynamic reorganization to obtain the network connectivity that may be needed to support task performance.

In prior studies, we reported that nodes of the visual-biased network exhibited robust resting-state connectivity with purported domain-general brain regions, while nodes of the auditory-biased network exhibited robust resting-state connectivity with purported language network regions (Tobyne et al., 2017; Noyce et al., 2022), but we had not explicitly mapped the supramodal regions in individual subjects. The current resting-state findings showing strong functional connectivity between frontal visual-biased and supramodal networks, add evidence in support of the proposed link between visual-biased and domain-general regions. Although the present study lacks the full variety of tasks to make a strong claim of domain-general processing for the supramodal WM regions in individual subjects, our supplementary analysis (Supplementary Figure 1, Supplementary Table 1) demonstrates that supramodal frontal regions largely overlapped with a core subset of the domain-general network (Assem et al., 2020). We suggest that the supramodal WM regions comprise an important subset of the domain-general network. In addition to the functional connectivity evidence, previous task activation data also revealed a similar asymmetry. Visual-biased ROIs were recruited in auditory WM tasks and thus exhibited a degree of supramodal WM, while auditory-biased ROIs showed little recruitment in visual WM tasks and thus showed little evidence for supramodal WM (Noyce et al., 2022; Tobyne et al., 2025). Thus, there may be a general asymmetry in activation and connectivity between frontal auditory and visual networks with the visual network maintaining preferential access to supramodal network resources, while auditory processing must recruit both networks to perform auditory WM tasks with high precision. We speculate that this cortical asymmetry may relate to the well-studied Colavita Visual Dominance effect (Colavita, 1974; Posner et al., 1976; Spence, 2009) in which the simultaneous presentation of visual and auditory stimuli leads to the extinction of the auditory stimulus. Several leading accounts of this visual prepotency effect argue that it reflects competitive attentional interactions that are biased toward vision (Fang et al., 2020; Posner et al., 1976; Spence et al., 2012). Additionally, behavioral studies have reported a tighter link for domain-general cognitive processing with visual working memory than for auditory working memory in adults (Bo et al., 2011) and children (Alhamdan et al., 2023).

Multiple aspects of analyses likely benefit from individual-specific analysis methods applied (Braga & Buckner, 2017; Gratton et al., 2018; Kong et al., 2021; Michalka et al., 2015). Functional parcellation of frontal cortex included many small regions with neighboring regions that often belonged to different functional categories (i.e., visual-biased, auditory-biased, or supramodal). Under such conditions, group-level definitions of ROIs will yield a very substantial loss of statistical power (Fedorenko et al., 2010; Tobyne et al., 2018); individual-specific methods are vital to the very definition of the ROIs. Importantly for our study, task-based functional connectivity analyses gain substantial power from individual-specific analyses as compared to group analyses (Gratton et al., 2018). It is also noteworthy that individual-specific analyses increase the power to predict behavioral measures from functional connectivity data (Kong et al., 2021). Nevertheless, it will be important to replicate the present findings with larger populations and additional auditory and visual tasks to more firmly demonstrate generality.

It should be noted that all functional scans required participants to maintain visual fixation and exposed them to auditory distractors due to the scanner noise. The requirement to maintain fixation was an important control to minimize neural activity related to eye movements and to equate eye position across all conditions, since eye movements can evoke substantial activity across the visual attentional network (Corbetta et al., 1998). The auditory noise, while minimized with hearing protections, is an unavoidable aspect of fMRI scanning. It is possible that one or both factors influenced the resting-state observations. Although there are prior reports of differences in functional connectivity between networks in eyes closed and fixation resting state conditions, the direction and networks involved vary considerably between studies (Agcaoglu et al., 2019; Han et al., 2023; Patriat et al., 2013). Because fixating on a single dot is a low-level visual task, if this task boosts visual connectivity to supramodal WM regions one might expect positive connectivity between lower-level visual cortical areas and supramodal regions, but the resting-state functional connectivity between *posterior visual* and *supramodal* networks was negative. Although we cannot conclusively rule out a small influence of the eyes-open fixation task and/or auditory scanner noise on the resting-state connectivity in frontal cortex, these conditions are constant and thus should not have affected the task-based gPPI analyses of changes in connectivity between task conditions (Figures 6, 7). We also acknowledge that the use of different scan parameters for task and resting-state analyses could potentially contribute to some difference between their results, although we note that the prior Noyce et al., 2022 report from our lab observed a similar resting-state network asymmetry using the same TR, SMS, and number of slices used here in the task study.

A final important caveat is that fMRI reports of dynamic network reorganization, including the present study, are potentially consistent with multiple interpretations at the level of individual neurons. Homogeneous populations of neurons within each ROI may alter their functional connectivity with neurons in other nodes and networks. Alternatively, ROIs may contain static, but heterogeneous neural populations that differ both in their connectivity and functional responses. Task activation of different sub-populations with different connectivity could appear as a dynamic network reconfiguration in the present type of analysis, even though individual neuronal connections are unchanging. Finer-scale analyses, possibly including those in non-human primates, are needed to distinguish between these alternatives.

This work uncovers several potentially fruitful pathways for further investigation. Future studies should examine network interactions under conditions where visual WM and auditory WM directly compete for resources and under conditions where visual and auditory features are integrated into multisensory object representations. Additional work should be conducted to better understand the representational content of each ROI in this analysis to elucidate their exact functional contribution. Other schemas for the functional organization of the frontal cortex could be added to this style of analysis, such as the language-specific frontal regions identified in Fedorenko et al. (2010, 2012). This would further integrate different lines of research into the functional organization of the frontal cortex. Lastly, further work relating behavioral measures to task-based changes in functional connectivity may have potential to be extended to clinically relevant dimensions of behavior.

This study builds on a body of work trying to make sense of the functional organization of the least clearly understood cortical region in the brain. Despite the lack of consensus on the functional organization of the frontal cortex, it is implicated in most all higher-level cognitive processes, (Amodio & Frith, 2006; Badre & Nee, 2018; De La Vega et al., 2016; Fuster, 2000) many neurological and psychiatric disorders, (Ferguson & Gao, 2018; Friedman & Robbins, 2022; Gao et al., 2012; Goldstein & Volkow, 2011; Goto et al., 2010) as well as the natural aging process (Giorgio et al., 2010; Nyberg et al., 2010). The results presented here drive the continued integration of multiple models of the functional organization of the frontal cortex, in particular, working to identify discrete functional networks for sensory-biased and supramodal processing and beginning to uncover how these networks interact during cognitive tasks and correlate with behavior.

## Data and Code Availability

All custom code and ROI search spaces (freesurfer label files in fsaverage space) used in these analyses are available at https://github.com/fmri/Sensory_Networks_FC. Data used to generate all figures are available in the supplementary materials. Data used to generate all figures are available in the supplemental materials.

## CRediT Author Statement

**Thomas Possidente**: Conceptualization, Formal analysis, Methodology, Software, Validation, Visualization, Writing - original draft, review & editing, **Vaibhav Tripathi**: Conceptualization, Data curation, Investigation, Methodology, Software, Writing – review & editing, **Joseph T. McGuire**: Methodology, Writing – review & editing **David Somers**: Conceptualization, Funding acquisition, Methodology, Supervision, Project administration, Writing - original draft, review & editing.

## Acknowledgments

This work is funded by National Science Foundation grant BCS-1829394 to D.C.S. This work involved the use of instrumentation supported by the NSF Major Research Instrumentation grant BCS-1625552. Data was analyzed on a high-performance computing cluster supported by the ONR grant N00014-17-1-2304.

We thank Dr. David Beeler for assistance with data collection and for helpful discussions, Dr. Abigail Noyce for auditory stimuli, and Ryan Marshall, Dr. Stephanie McMains, and Shruthi Chakrapani for scanning assistance. We acknowledge the University of Minnesota Center for Magnetic Resonance Research for use of the multiband-EPI pulse sequences.

## References

Agcaoglu, O., Wilson, T. W., Wang, Y.-P., Stephen, J., & Calhoun, V. D. (2019). Resting state connectivity differences in eyes open versus eyes closed conditions. Human Brain Mapping, 40(8), 2488–2498. 10.1002/hbm.24539

Alhamdan, A. A., Murphy, M. J., Pickering, H. E., & Crewther, S. G. (2023). The Contribution of Visual and Auditory Working Memory and Non-Verbal IQ to Motor Multisensory Processing in Elementary School Children. Brain Sciences, 13(2), Article 2. 10.3390/brainsci13020270

Alvarez, G. A., & Cavanagh, P. (2004). The capacity of visual short-term memory is set both by visual information load and by number of objects. Psychological Science : A Journal of the American Psychological Society / APS, 15(2), 106–111.

Amodio, D. M., & Frith, C. D. (2006). Meeting of minds: The medial frontal cortex and social cognition. Nature Reviews Neuroscience, 7(4), 268–277. 10.1038/nrn1884

Ashburner, J. (2007). A fast diffeomorphic image registration algorithm. NeuroImage, 38(1), 95–113. 10.1016/j.neuroimage.2007.07.007

Ashburner, J., & Friston, K. (1997). Multimodal Image Coregistration and Partitioning—A Unified Framework. NeuroImage, 6(3), 209–217. 10.1006/nimg.1997.0290

Ashburner, J., & Friston, K. J. (2005). Unified segmentation. NeuroImage, 26(3), 839–851. 10.1016/j.neuroimage.2005.02.018

Assem, M., Glasser, M. F., Van Essen, D. C., & Duncan, J. (2020). A Domain-General Cognitive Core Defined in Multimodally Parcellated Human Cortex. Cerebral Cortex, 30(8), 4361–4380. 10.1093/cercor/bhaa023

Baddeley, A. D., & Hitch, G. (1974). Working Memory. In G. H. Bower (Ed.), Psychology of Learning and Motivation (Vol. 8, pp. 47–89). Academic Press. 10.1016/S0079-7421(08)60452-1

Badre, D., & Nee, D. E. (2018). Frontal Cortex and the Hierarchical Control of Behavior. Trends in Cognitive Sciences, 22(2), 170–188. 10.1016/j.tics.2017.11.005

Bar-Joseph, Z., Gifford, D. K., & Jaakkola, T. S. (2001). Fast optimal leaf ordering for hierarchical clustering. Bioinformatics, 17(suppl_1), S22–S29. 10.1093/bioinformatics/17.suppl_1.S22

Barr, D. J., Levy, R., Scheepers, C., & Tily, H. J. (2013). Random effects structure for confirmatory hypothesis testing: Keep it maximal. Journal of Memory and Language, 68(3), 255–278. 10.1016/j.jml.2012.11.001

Bays, P. M., Catalao, R. F. G., & Husain, M. (2009). The precision of visual working memory is set by allocation of a shared resource. Journal of Vision, 9(10), 7. 10.1167/9.10.7

Behzadi, Y., Restom, K., Liau, J., & Liu, T. T. (2007). A component based noise correction method (CompCor) for BOLD and perfusion based fMRI. NeuroImage, 37(1), 90–101. 10.1016/j.neuroimage.2007.04.042

Bettencourt, K. C., & Xu, Y. (2016). Decoding the content of visual short-term memory under distraction in occipital and parietal areas. Nature Neuroscience, 19(1), 150–157. 10.1038/nn.4174

Bo, J., Jennett, S., & Seidler, R. D. (2011). Working memory capacity correlates with implicit serial reaction time task performance. Experimental Brain Research, 214(1), 73–81. 10.1007/s00221-011-2807-8

Braga, R. M., & Buckner, R. L. (2017). Parallel Interdigitated Distributed Networks within the Individual Estimated by Intrinsic Functional Connectivity. Neuron, 95(2), 457–471.e5. 10.1016/j.neuron.2017.06.038

Cauley, S. F., Polimeni, J. R., Bhat, H., Wald, L. L., & Setsompop, K. (2014). Interslice leakage artifact reduction technique for simultaneous multislice acquisitions. Magnetic Resonance in Medicine, 72(1), 93–102. 10.1002/mrm.24898

Chai, X. J., Castañón, A. N., Öngür, D., & Whitfield-Gabrieli, S. (2012). Anticorrelations in resting state networks without global signal regression. NeuroImage, 59(2), 1420–1428. 10.1016/j.neuroimage.2011.08.048

Christophel, T. B., Hebart, M. N., & Haynes, J.-D. (2012). Decoding the Contents of Visual Short-Term Memory from Human Visual and Parietal Cortex. Journal of Neuroscience, 32(38), 12983–12989. 10.1523/JNEUROSCI.0184-12.2012

Christophel, T. B., Klink, P. C., Spitzer, B., Roelfsema, P. R., & Haynes, J.-D. (2017). The Distributed Nature of Working Memory. Trends in Cognitive Sciences, 21(2), 111–124. 10.1016/j.tics.2016.12.007

Cisler, J. M., Bush, K., & Steele, J. S. (2014). A comparison of statistical methods for detecting context-modulated functional connectivity in fMRI. NeuroImage, 84, 1042–1052. 10.1016/j.neuroimage.2013.09.018

Colavita, F. B. (1974). Human sensory dominance. Perception & Psychophysics, 16(2), 409–412. 10.3758/BF03203962

Cole, M. W., Ito, T., Cocuzza, C., & Sanchez-Romero, R. (2021). The Functional Relevance of Task-State Functional Connectivity. The Journal of Neuroscience, 41(12), 2684–2702. 10.1523/JNEUROSCI.1713-20.2021

Corbetta, M., Akbudak, E., Conturo, T. E., Snyder, A. Z., Ollinger, J. M., Drury, H. A., Linenweber, M. R., Petersen, S. E., Raichle, M. E., Van Essen, D. C., & Shulman, G. L. (1998). A common network of functional areas for attention and eye movements. Neuron, 21(4), 761–773.

Crittenden, B. M., & Duncan, J. (2014). Task Difficulty Manipulation Reveals Multiple Demand Activity but no Frontal Lobe Hierarchy. Cerebral Cortex, 24(2), 532–540. 10.1093/cercor/bhs333

Dale, A. M., Fischl, B., & Sereno, M. I. (1999). Cortical Surface-Based Analysis: I. Segmentation and Surface Reconstruction. NeuroImage, 9(2), 179–194. 10.1006/nimg.1998.0395

De La Vega, A., Chang, L. J., Banich, M. T., Wager, T. D., & Yarkoni, T. (2016). Large-Scale Meta-Analysis of Human Medial Frontal Cortex Reveals Tripartite Functional Organization. The Journal of Neuroscience, 36(24), 6553–6562. 10.1523/JNEUROSCI.4402-15.2016

D’Esposito, M., & Postle, B. R. (2015). The Cognitive Neuroscience of Working Memory. Annual Review of Psychology, 66(Volume 66, 2015), 115–142. 10.1146/annurev-psych-010814-015031

Dice, L. R. (1945). Measures of the Amount of Ecologic Association Between Species. Ecology, 26(3), 297–302. 10.2307/1932409

Duncan, J., & Owen, A. M. (2000). Common regions of the human frontal lobe recruited by diverse cognitive demands. Trends in Neurosciences, 23(10), 475–483. 10.1016/S0166-2236(00)01633-7

Eklund, A., Knutsson, H., & Nichols, T. E. (2019). Cluster failure revisited: Impact of first level design and physiological noise on cluster false positive rates. Human Brain Mapping, 40(7), 2017–2032. 10.1002/hbm.24350

Eklund, A., Nichols, T. E., & Knutsson, H. (2016). Cluster failure: Why fMRI inferences for spatial extent have inflated false-positive rates. Proceedings of the National Academy of Sciences, 113(28), 7900–7905. 10.1073/pnas.1602413113

Ester, E. F., Sprague, T. C., & Serences, J. T. (2015). Parietal and Frontal Cortex Encode Stimulus-Specific Mnemonic Representations during Visual Working Memory. Neuron, 87(4), 893–905. 10.1016/j.neuron.2015.07.013

Fang, Y., Li, Y., Xu, X., Tao, H., & Chen, Q. (2020). Top-down attention modulates the direction and magnitude of sensory dominance. Experimental Brain Research, 238(3), 587–600. 10.1007/s00221-020-05737-7

Fedorenko, E., Duncan, J., & Kanwisher, N. (2012). Language-Selective and Domain-General Regions Lie Side by Side within Broca’s Area. Current Biology, 22(21), 2059–2062. 10.1016/j.cub.2012.09.011

Fedorenko, E., Hsieh, P.-J., Nieto-Castañón, A., Whitfield-Gabrieli, S., & Kanwisher, N. (2010). New Method for fMRI Investigations of Language: Defining ROIs Functionally in Individual Subjects. Journal of Neurophysiology, 104(2), 1177–1194. 10.1152/jn.00032.2010

Feinberg, D. A., Moeller, S., Smith, S. M., Auerbach, E., Ramanna, S., Glasser, M. F., Miller, K. L., Ugurbil, K., & Yacoub, E. (2010). Multiplexed Echo Planar Imaging for Sub-Second Whole Brain FMRI and Fast Diffusion Imaging. PLOS ONE, 5(12), e15710. 10.1371/journal.pone.0015710

Ferguson, B. R., & Gao, W.-J. (2018). PV Interneurons: Critical Regulators of E/I Balance for Prefrontal Cortex-Dependent Behavior and Psychiatric Disorders. Frontiers in Neural Circuits, 12. 10.3389/fncir.2018.00037

Fischl, B. (2012). FreeSurfer. NeuroImage, 62(2), 774–781. 10.1016/j.neuroimage.2012.01.021

Fischl, B., Liu, A., & Dale, A. M. (2001). Automated manifold surgery: Constructing geometrically accurate and topologically correct models of the human cerebral cortex. IEEE Transactions on Medical Imaging, 20(1), 70–80. IEEE Transactions on Medical Imaging. 10.1109/42.906426

Fischl, B., Salat, D. H., Busa, E., Albert, M., Dieterich, M., Haselgrove, C., Kouwe, A. van der, Killiany, R., Kennedy, D., Klaveness, S., Montillo, A., Makris, N., Rosen, B., & Dale, A. M. (2002). Whole Brain Segmentation: Automated Labeling of Neuroanatomical Structures in the Human Brain. Neuron, 33(3), 341–355. 10.1016/S0896-6273(02)00569-X

Franconeri, S. L., Alvarez, G. A., & Cavanagh, P. (2013). Flexible cognitive resources: Competitive content maps for attention and memory. Trends in Cognitive Sciences, 17(3), 134–141. 10.1016/j.tics.2013.01.010

Friedman, N. P., & Robbins, T. W. (2022). The role of prefrontal cortex in cognitive control and executive function. Neuropsychopharmacology, 47(1), 72–89. 10.1038/s41386-021-01132-0

Friston, K. J., Buechel, C., Fink, G. R., Morris, J., Rolls, E., & Dolan, R. J. (1997). Psychophysiological and Modulatory Interactions in Neuroimaging. NeuroImage, 6(3), 218–229. 10.1006/nimg.1997.0291

Fuster, J. M. (2000). Executive frontal functions. Experimental Brain Research, 133(1), 66–70. 10.1007/s002210000401

Gao, W.-J., Wang, H.-X., A., M., & Li, Y.-C. (2012). The Unique Properties of the Prefrontal Cortex and Mental Illness. In T. Mantamadiotis (Ed.), When Things Go Wrong—Diseases and Disorders of the Human Brain. InTech. 10.5772/35868

Giorgio, A., Santelli, L., Tomassini, V., Bosnell, R., Smith, S., De Stefano, N., & Johansen-Berg, H. (2010). Age-related changes in grey and white matter structure throughout adulthood. NeuroImage, 51(3), 943–951. 10.1016/j.neuroimage.2010.03.004

Glasser, M. F., Coalson, T. S., Robinson, E. C., Hacker, C. D., Harwell, J., Yacoub, E., Ugurbil, K., Andersson, J., Beckmann, C. F., Jenkinson, M., Smith, S. M., & Van Essen, D. C. (2016). A multi-modal parcellation of human cerebral cortex. Nature, 536(7615), Article 7615. 10.1038/nature18933

Goldstein, R. Z., & Volkow, N. D. (2011). Dysfunction of the prefrontal cortex in addiction: Neuroimaging findings and clinical implications. Nature Reviews Neuroscience, 12(11), 652–669. 10.1038/nrn3119

Goto, Y., Yang, C. R., & Otani, S. (2010). Functional and Dysfunctional Synaptic Plasticity in Prefrontal Cortex: Roles in Psychiatric Disorders. Biological Psychiatry, 67(3), 199–207. 10.1016/j.biopsych.2009.08.026

Gratton, C., Laumann, T. O., Nielsen, A. N., Greene, D. J., Gordon, E. M., Gilmore, A. W., Nelson, S. M., Coalson, R. S., Snyder, A. Z., Schlaggar, B. L., Dosenbach, N. U. F., & Petersen, S. E. (2018). Functional Brain Networks Are Dominated by Stable Group and Individual Factors, Not Cognitive or Daily Variation. Neuron, 98(2), 439–452.e5. 10.1016/j.neuron.2018.03.035

Griswold, M. A., Jakob, P. M., Heidemann, R. M., Nittka, M., Jellus, V., Wang, J., Kiefer, B., & Haase, A. (2002). Generalized autocalibrating partially parallel acquisitions (GRAPPA). Magnetic Resonance in Medicine, 47(6), 1202–1210. 10.1002/mrm.10171

Hagler, D. J., Saygin, A. P., & Sereno, M. I. (2006). Smoothing and cluster thresholding for cortical surface-based group analysis of fMRI data. NeuroImage, 33(4), 1093–1103. 10.1016/j.neuroimage.2006.07.036

Han, J., Zhou, L., Wu, H., Huang, Y., Qiu, M., Huang, L., Lee, C., Lane, T. J., & Qin, P. (2023). Eyes-Open and Eyes-Closed Resting State Network Connectivity Differences. Brain Sciences, 13(1), Article 1. 10.3390/brainsci13010122

Harrison, S. A., & Tong, F. (2009). Decoding reveals the contents of visual working memory in early visual areas. Nature, 458(7238), 632–635. 10.1038/nature07832

Hartcher-O’Brien, J., Gallace, A., Krings, B., Koppen, C., & Spence, C. (2008). When vision ‘extinguishes’ touch in neurologically-normal people: Extending the Colavita visual dominance effect. Experimental Brain Research, 186(4), 643–658. 10.1007/s00221-008-1272-5

Hartman, J. (2023). Emmeans [MATLAB]. https://github.com/jackatta/estimated-marginal-means (Original work published 2019)

Hecht, D., & Reiner, M. (2009). Sensory dominance in combinations of audio, visual and haptic stimuli. Experimental Brain Research, 193(2), 307–314. 10.1007/s00221-008-1626-z

Hedges, L. V. (2007). Effect Sizes in Cluster-Randomized Designs. Journal of Educational and Behavioral Statistics, 32(4), 341–370. 10.3102/1076998606298043

Jenkinson, M., Beckmann, C. F., Behrens, T. E. J., Woolrich, M. W., & Smith, S. M. (2012). FSL. NeuroImage, 62(2), 782–790. 10.1016/j.neuroimage.2011.09.015

Kong, R., Yang, Q., Gordon, E., Xue, A., Yan, X., Orban, C., Zuo, X.-N., Spreng, N., Ge, T., Holmes, A., Eickhoff, S., & Yeo, B. T. T. (2021). Individual-Specific Areal-Level Parcellations Improve Functional Connectivity Prediction of Behavior. Cerebral Cortex, 31(10), 4477–4500. 10.1093/cercor/bhab101

Krienen, F. M., Yeo, B. T. T., & Buckner, R. L. (2014). Reconfigurable task-dependent functional coupling modes cluster around a core functional architecture. Philosophical Transactions of the Royal Society of London. Series B, Biological Sciences, 369(1653), 20130526. 10.1098/rstb.2013.0526

Linke, A. C., Vicente-Grabovetsky, A., & Cusack, R. (2011). Stimulus-specific suppression preserves information in auditory short-term memory. Proceedings of the National Academy of Sciences, 108(31), 12961–12966. 10.1073/pnas.1102118108

Mayer, A. R., Ryman, S. G., Hanlon, F. M., Dodd, A. B., & Ling, J. M. (2017). Look Hear! The Prefrontal Cortex is Stratified by Modality of Sensory Input During Multisensory Cognitive Control. Cerebral Cortex, 27(5), 2831–2840. 10.1093/cercor/bhw131

McLaren, D. G., Ries, M. L., Xu, G., & Johnson, S. C. (2012). A generalized form of context-dependent psychophysiological interactions (gPPI): A comparison to standard approaches. NeuroImage, 61(4), 1277–1286. 10.1016/j.neuroimage.2012.03.068

Michalka, S. W., Kong, L., Rosen, M. L., Shinn-Cunningham, B. G., & Somers, D. C. (2015). Short-Term Memory for Space and Time Flexibly Recruit Complementary Sensory-Biased Frontal Lobe Attention Networks. Neuron, 87(4), 882–892. 10.1016/j.neuron.2015.07.028

Moeller, S., Yacoub, E., Olman, C. A., Auerbach, E., Strupp, J., Harel, N., & Uğurbil, K. (2010). Multiband multislice GE-EPI at 7 tesla, with 16-fold acceleration using partial parallel imaging with application to high spatial and temporal whole-brain fMRI. Magnetic Resonance in Medicine, 63(5), 1144–1153. 10.1002/mrm.22361

Nieto-Castanon, A. (2020). Handbook of functional connectivity Magnetic Resonance Imaging methods in CONN. Hilbert Press.

Noyce, A. L., Cestero, N., Michalka, S. W., Shinn-Cunningham, B. G., & Somers, D. C. (2017). Sensory-Biased and Multiple-Demand Processing in Human Lateral Frontal Cortex. Journal of Neuroscience, 37(36), 8755–8766. 10.1523/JNEUROSCI.0660-17.2017

Noyce, A. L., Lefco, R. W., Brissenden, J. A., Tobyne, S. M., Shinn-Cunningham, B. G., & Somers, D. C. (2022). Extended Frontal Networks for Visual and Auditory Working Memory. Cerebral Cortex, 32(4), 855–869. 10.1093/cercor/bhab249

Nyberg, L., Salami, A., Andersson, M., Eriksson, J., Kalpouzos, G., Kauppi, K., Lind, J., Pudas, S., Persson, J., & Nilsson, L.-G. (2010). Longitudinal evidence for diminished frontal cortex function in aging. Proceedings of the National Academy of Sciences, 107(52), 22682–22686. 10.1073/pnas.1012651108

O’Reilly, J. X., Woolrich, M. W., Behrens, T. E. J., Smith, S. M., & Johansen-Berg, H. (2012). Tools of the trade: Psychophysiological interactions and functional connectivity. Social Cognitive and Affective Neuroscience, 7(5), 604–609. 10.1093/scan/nss055

Patriat, R., Molloy, E. K., Meier, T. B., Kirk, G. R., Nair, V. A., Meyerand, M. E., Prabhakaran, V., & Birn, R. M. (2013). The effect of resting condition on resting-state fMRI reliability and consistency: A comparison between resting with eyes open, closed, and fixated. NeuroImage, 78, 463–473. 10.1016/j.neuroimage.2013.04.013

Penny, W. D., Friston, K. J., Ashburner, J. T., Kiebel, S. J., & Nichols, T. E. (2011). Statistical Parametric Mapping: The Analysis of Functional Brain Images. Elsevier.

Posner, M. I., Nissen, M. J., & Klein, R. M. (1976). Visual dominance: An information-processing account of its origins and significance. Psychological Review, 83(2), 157–171. 10.1037/0033-295X.83.2.157

Power, J. D., Mitra, A., Laumann, T. O., Snyder, A. Z., Schlaggar, B. L., & Petersen, S. E. (2014). Methods to detect, characterize, and remove motion artifact in resting state fMRI. NeuroImage, 84, 320–341. 10.1016/j.neuroimage.2013.08.048

Ségonne, F., Dale, A. M., Busa, E., Glessner, M., Salat, D., Hahn, H. K., & Fischl, B. (2004). A hybrid approach to the skull stripping problem in MRI. NeuroImage, 22(3), 1060–1075. 10.1016/j.neuroimage.2004.03.032

Segonne, F., Pacheco, J., & Fischl, B. (2007). Geometrically Accurate Topology-Correction of Cortical Surfaces Using Nonseparating Loops. IEEE Transactions on Medical Imaging, 26(4), 518–529. IEEE Transactions on Medical Imaging. 10.1109/TMI.2006.887364

Serences, J. T., Ester, E. F., Vogel, E. K., & Awh, E. (2009). Stimulus-Specific Delay Activity in Human Primary Visual Cortex. Psychological Science, 20(2), 207–214. 10.1111/j.1467-9280.2009.02276.x

Setsompop, K., Cohen-Adad, J., Gagoski, B. A., Raij, T., Yendiki, A., Keil, B., Wedeen, V. J., & Wald, L. L. (2012). Improving diffusion MRI using simultaneous multi-slice echo planar imaging. NeuroImage, 63(1), 569–580. 10.1016/j.neuroimage.2012.06.033

Sladky, R., Friston, K. J., Tröstl, J., Cunnington, R., Moser, E., & Windischberger, C. (2011). Slice-timing effects and their correction in functional MRI. NeuroImage, 58(2), 588–594. 10.1016/j.neuroimage.2011.06.078

Sled, J. G., Zijdenbos, A. P., & Evans, A. C. (1998). A nonparametric method for automatic correction of intensity nonuniformity in MRI data. IEEE Transactions on Medical Imaging, 17(1), 87–97. IEEE Transactions on Medical Imaging. 10.1109/42.668698

Smith, S. M., Fox, P. T., Miller, K. L., Glahn, D. C., Fox, P. M., Mackay, C. E., Filippini, N., Watkins, K. E., Toro, R., Laird, A. R., & Beckmann, C. F. (2009). Correspondence of the brain’s functional architecture during activation and rest. Proceedings of the National Academy of Sciences, 106(31), 13040–13045. 10.1073/pnas.0905267106

Smith, S. M., Jenkinson, M., Woolrich, M. W., Beckmann, C. F., Behrens, T. E. J., Johansen-Berg, H., Bannister, P. R., De Luca, M., Drobnjak, I., Flitney, D. E., Niazy, R. K., Saunders, J., Vickers, J., Zhang, Y., De Stefano, N., Brady, J. M., & Matthews, P. M. (2004). Advances in functional and structural MR image analysis and implementation as FSL. NeuroImage, 23, S208–S219. 10.1016/j.neuroimage.2004.07.051

Smith, S. M., & Nichols, T. E. (2009). Threshold-free cluster enhancement: Addressing problems of smoothing, threshold dependence and localisation in cluster inference. NeuroImage, 44(1), 83–98. 10.1016/j.neuroimage.2008.03.061

Sørensen, T. (1948). A method of establishing groups of equal amplitude in plant sociology based on similarity of species and its application to analyses of the vegetation on Danish commons. In Biol Skrifter (Vol. 5, p. 1). https://cir.nii.ac.jp/crid/1370302864784225926

Spence, C. (2009). Explaining the Colavita visual dominance effect. In N. Srinivasan (Ed.), Progress in Brain Research (Vol. 176, pp. 245–258). Elsevier. 10.1016/S0079-6123(09)17615-X

Spence, C., Parise, C., & Chen, Y.-C. (2012). The Colavita Visual Dominance Effect. In M. M. Murray & M. T. Wallace (Eds.), The Neural Bases of Multisensory Processes. CRC Press/Taylor & Francis. http://www.ncbi.nlm.nih.gov/books/NBK92851/

Studholme, C., Hawkes, D. J., & Hill, D. L. G. (1998). Normalized entropy measure for multimodality image alignment. Medical Imaging 1998: Image Processing, 3338, 132–143. 10.1117/12.310835

Tobyne, S. M., Brissenden, J. A., Noyce, A. L., & Somers, D. C. (2025). Combined Auditory, Tactile, and Visual FMRI Reveals Sensory-Biased and Supramodal Working Memory Regions in the Human Frontal Cortex. Journal of Neuroscience, 45(38). 10.1523/JNEUROSCI.0773-25.2025

Tobyne, S. M., Osher, D. E., Michalka, S. W., & Somers, D. C. (2017). Sensory-biased attention networks in human lateral frontal cortex revealed by intrinsic functional connectivity. NeuroImage, 162, 362–372. 10.1016/j.neuroimage.2017.08.020

Tobyne, S. M., Somers, D. C., Brissenden, J. A., Michalka, S. W., Noyce, A. L., & Osher, D. E. (2018). Prediction of individualized task activation in sensory modality-selective frontal cortex with ‘connectome fingerprinting.’ NeuroImage, 183, 173–185. 10.1016/j.neuroimage.2018.08.007

Uluç, I., Schmidt, T. T., Wu, Y., & Blankenburg, F. (2018). Content-specific codes of parametric auditory working memory in humans. NeuroImage, 183, 254–262. 10.1016/j.neuroimage.2018.08.024

van der Kouwe, A. J. W., Benner, T., Salat, D. H., & Fischl, B. (2008). Brain morphometry with multiecho MPRAGE. NeuroImage, 40(2), 559–569. 10.1016/j.neuroimage.2007.12.025

Whitfield-Gabrieli, S., & Nieto-Castanon, A. (2011). Artifact Detection Tools (ART). https://www.nitrc.org/projects/artifact_detect

Whitfield-Gabrieli, S., & Nieto-Castanon, A. (2012). Conn: A Functional Connectivity Toolbox for Correlated and Anticorrelated Brain Networks. Brain Connectivity, 2(3), 125–141. 10.1089/brain.2012.0073

Xu, J., Moeller, S., Auerbach, E. J., Strupp, J., Smith, S. M., Feinberg, D. A., Yacoub, E., & Uğurbil, K. (2013). Evaluation of slice accelerations using multiband echo planar imaging at 3 T. NeuroImage, 83, 991–1001. 10.1016/j.neuroimage.2013.07.055

