## Supplemental Figures, Tables, Notes for "Interactions between sensory-biased and supramodal working memory networks in the human cerebral cortex"

### Supplementary Note 1: Resting-State Network Controls

To ensure that the results were not unexpectedly biased by the inclusion of individual ROIs defined with guidance from the probabilistic maps, we repeated the above LME analysis in full using only the ~80% of ROIs defined robustly from individual subject data alone. The connection between *frontal visual ↔ frontal+posterior auditory* networks dropped below significance in this alternative analysis, further indicating relatively segregated processing. All other significant and non-significant results remained (Supplementary Table 4), including the significant connectivity difference between *frontal visual ↔ supramodal* (M=.18, SE=.03) and the *frontal auditory ↔ supramodal* networks (M=.05, SE=.02) ($W(1)=4.7$, $p<.001$).

Additionally, to show that the overall results were not idiosyncratic to small changes in network clustering, we repeated the above LME analysis using a network grouping in which the *frontal+posterior auditory* network was replaced by separate *frontal auditory* and *posterior auditory* networks, to mirror the visual network structure. Also, midIFS was included in the *frontal visual* network, based on expected functionality (Noyce et al., 2022; Tobyne et al., 2025). The resulting connectivity structure was very similar (Supplementary Figure 5 and Supplementary Table 5), and critically, the connectivity between *frontal visual ↔ supramodal* (M=.19, SE=.02) was significantly greater ($W(1)=5.0$, $p<.001$) than between the *frontal auditory ↔ supramodal* networks (M=.07, SE=.02).

To address concerns about distance-dependent correlations, we included ROI-to-ROI distance as a random-effect in our LME analysis and turned off the modest 3mm smoothing kernel. We calculated distance between ROIs by first calculating the "medoid" for each ROI (vertex with smallest sum of Euclidean distances to all other vertices), then calculating the Euclidean distance between each pair of ROI medoids. With that change in the LME model, we still find a significant difference ($W(1)=3.30$, $p<.001$) in connectivity between *frontal visual ↔ supramodal* (M=.15, SE=.03) and the *frontal+posterior auditory ↔ supramodal* networks (M=.06, SE=.02).

**Supplementary Note 2: LME Assumption Checks**

In terms of distributional assumptions, the residuals of the LME models are not expected to pass statistical tests for normality because of the extreme sensitivity of these tests under high numbers of data points (>7k points derived from ipsilateral left, ipsilateral right, and contralateral connections between 16 ROIs in 21 individual participants). Linear regression models are robust to non-extreme deviations from normality (Knief & Forstmeier, 2021; Schmidt & Finan, 2018), and our connectivity and gPPI LME models display non-extreme skewness (.49, -.13 respectively) and excess kurtosis (1.46, 1.60 respectively) in their residuals. Additionally, the results from the non-parametric ROI-ROI TFCE analyses mirror those found in the LME analyses.

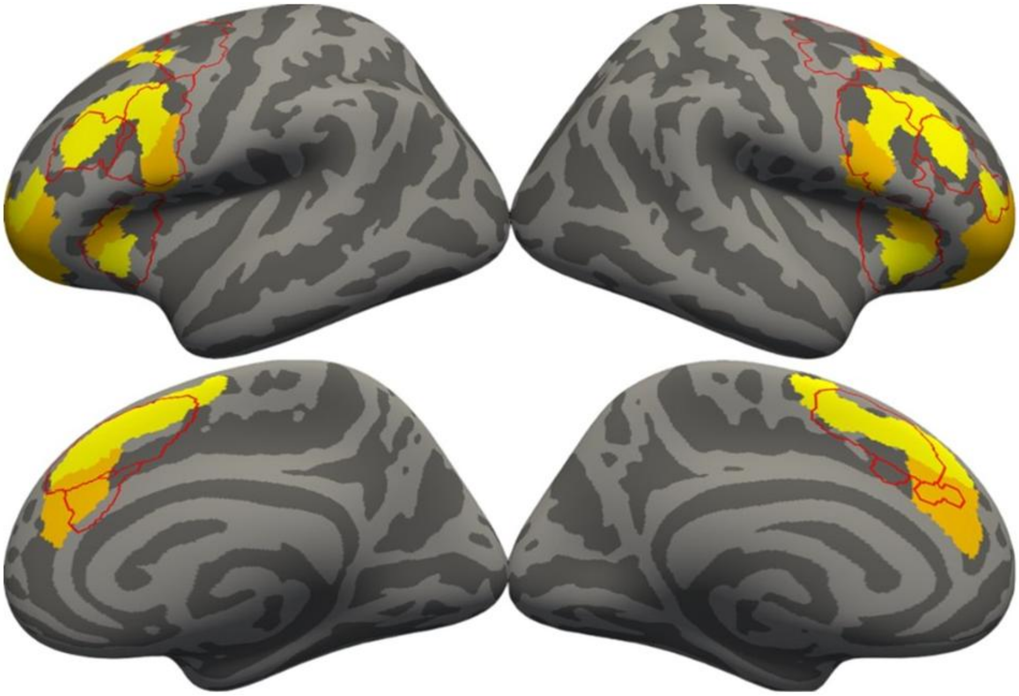

Supplementary Figure 1. Shows the overlap between supramodal ROI search spaces from this study (red outlines) and frontal domain-general ROIs identified in Assem et al. (2020) using WM, math/language, and relational reasoning tasks. Yellow ROIs represent “core” domain-general ROIs, while orange ROIs represent “penumbra” domain-general ROIs. Domain-general ROIs were displayed using the HCP-MMP1.0 parcellation projected onto fsaverage.

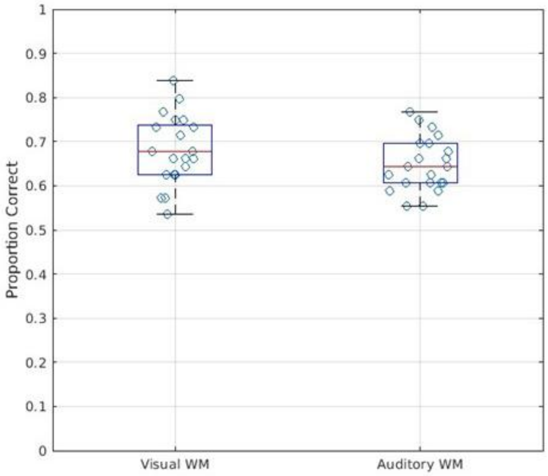


Supplementary Figure 2. Box plots of percentage correct responses for visual and auditory WM tasks overlayed with individual participant percent correct rate jittered scatter plots.

A)

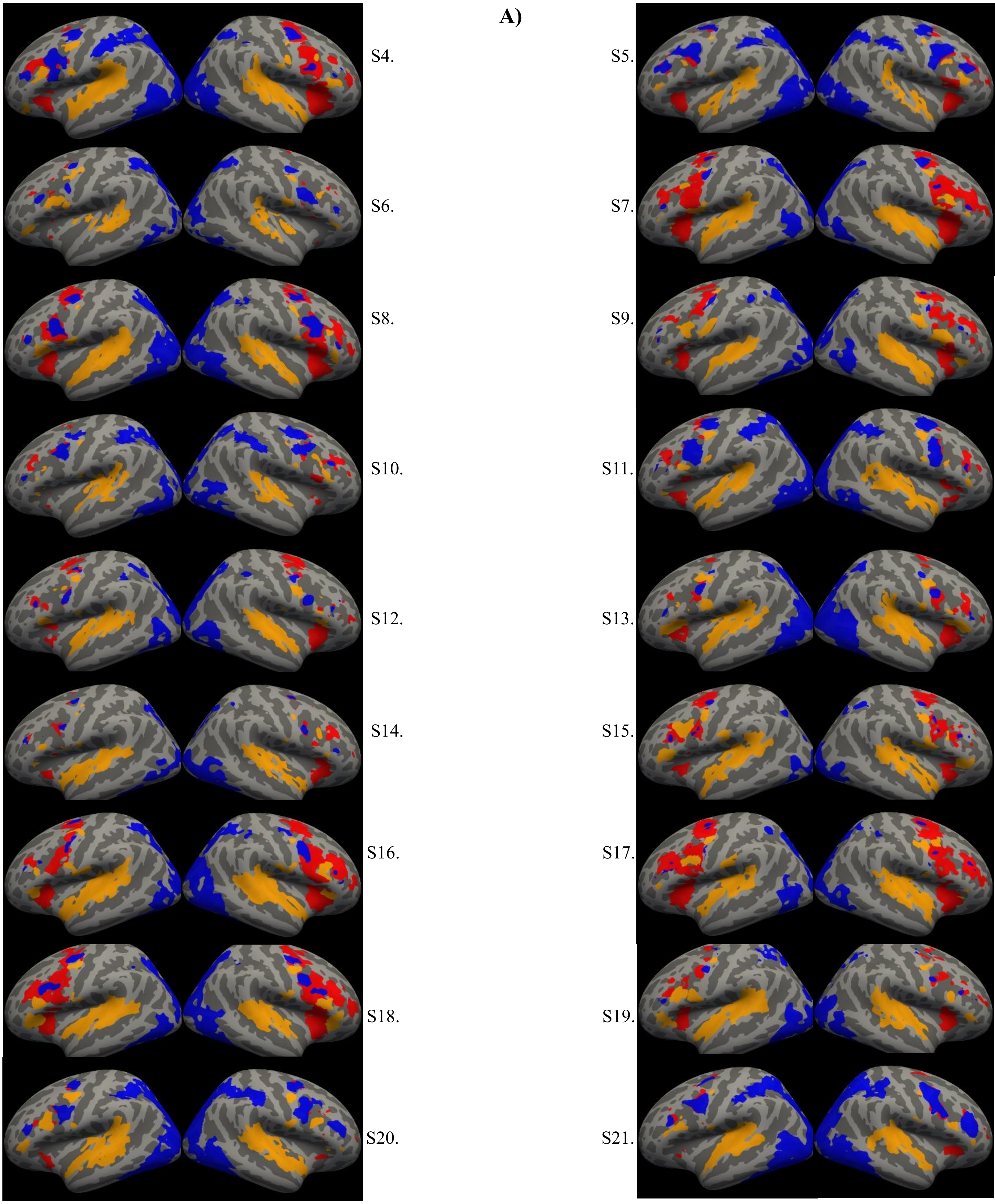

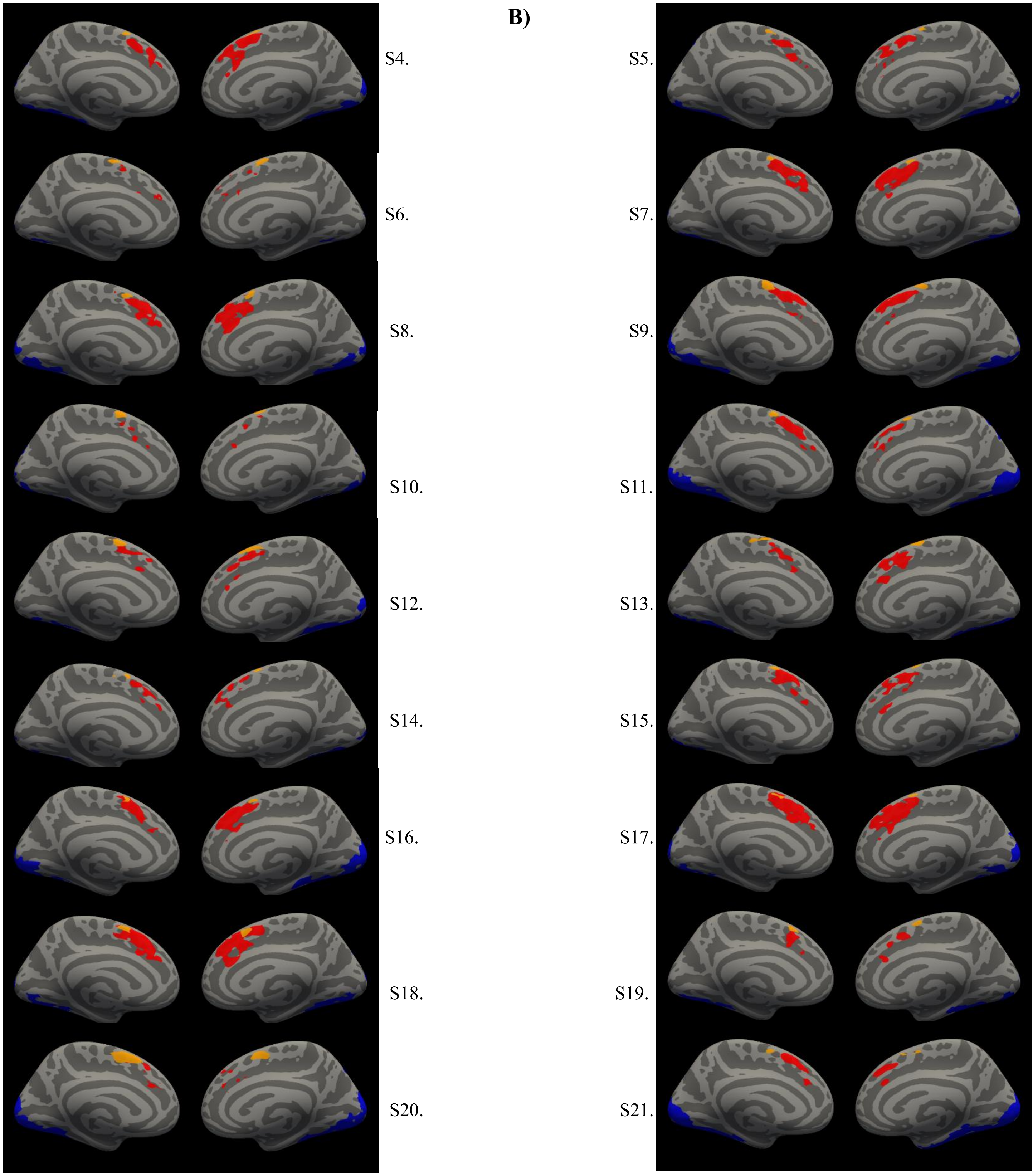


Supplementary Figure 3. ROIs for all subjects not presented in main text Figure 4. Blue areas are visual-biased, orange areas are auditory-biased, and red areas are supramodal A) Lateral view. B) Medial view.

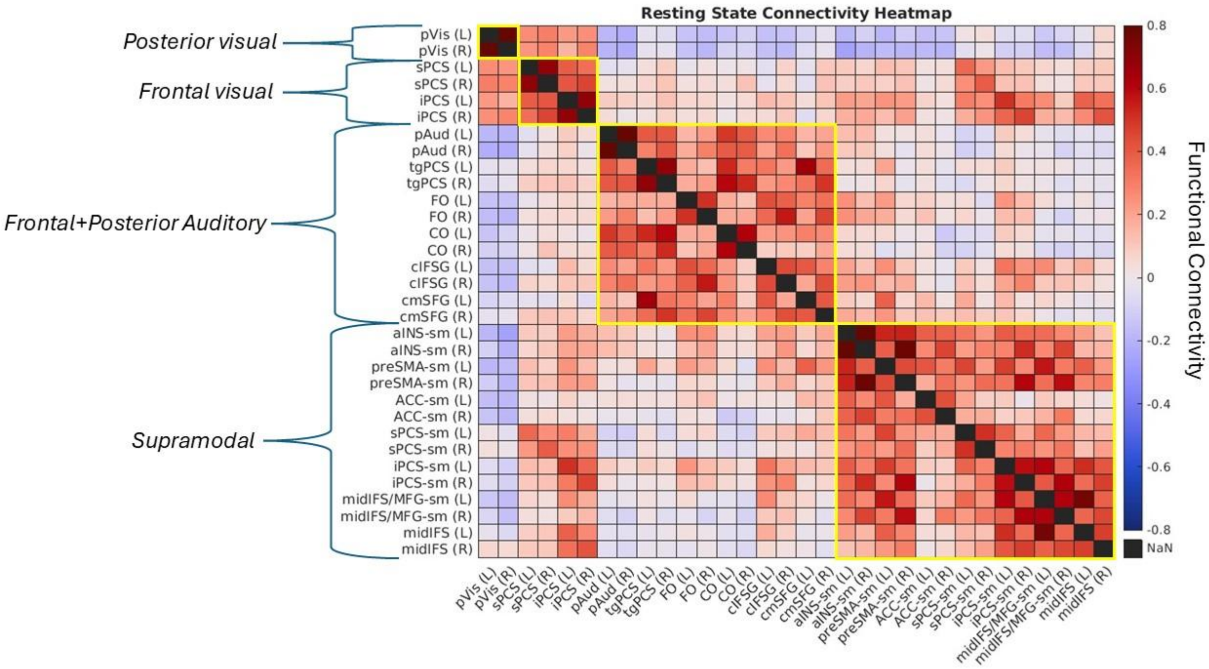


Supplementary Figure 4. Resting state functional connectivity matrix heatmap. Yellow borders represent HCA clusters. Functional connectivity values are from Conn Toolbox weighted-GLM derived connectivity coefficients.

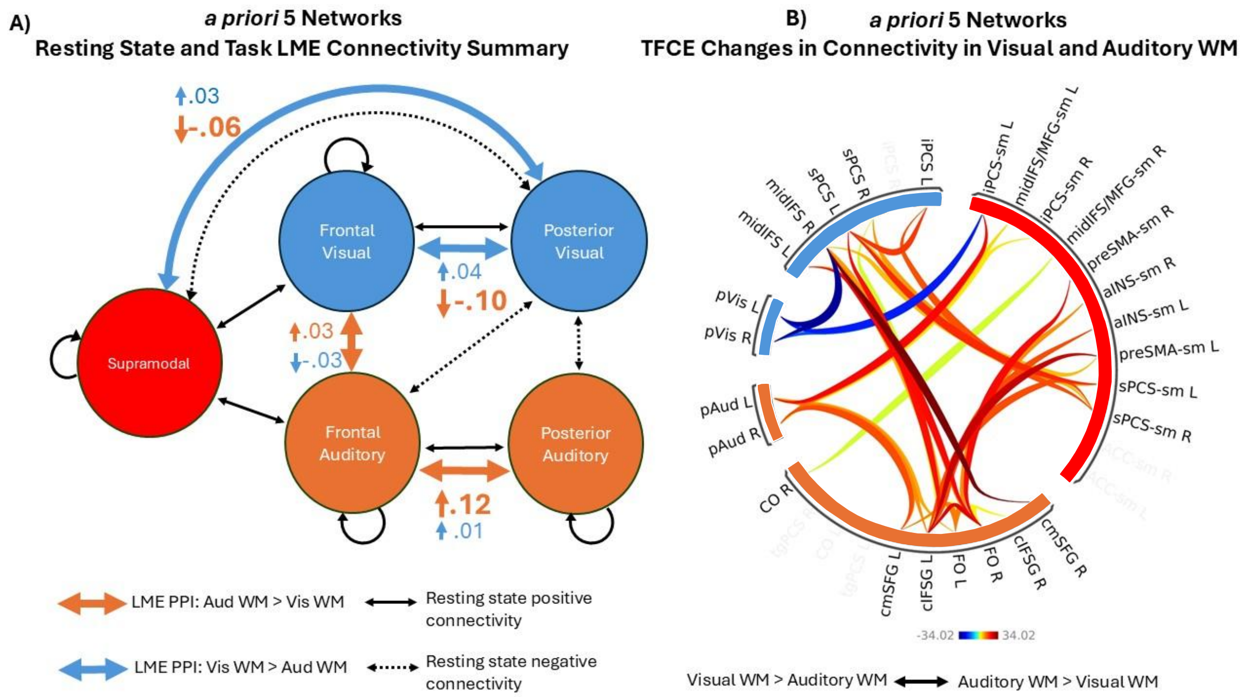


Supplementary Figure 5. Task-based functional connectivity results for the *a priori* symmetric network clustering. A) Diagram summarizing the LME-derived significant changes in connectivity between ROI networks during visual and auditory WM. The black arrows show significant resting-state connectivity, while the large colored arrows indicate change in connectivity. The values and small colored arrows shown next to the large colored arrows indicate the mean increase or decrease in PPI coefficients for the visual and auditory WM conditions between ROIs in each network. For example, the change in connectivity between *posterior visual* ←→ *frontal visual* networks increased under visual WM (large blue arrow) and was driven by a small (.04) increase in the visual WM condition and a large decrease (.10) in the auditory condition (small blue and orange arrows and values). Changes in self-connection in the *posterior visual* and *auditory* networks were not assessed because the network only contained 1 (bilateral) ROI. C) Connectogram summarizing the TFCE-derived significant changes in connectivity between individual ROIs during visual and auditory WM. Warm colors represent auditory>visual changes in connectivity while cool colors represent visual>auditory changes in connectivity. ROIs are grouped around the circle by HCA-derived networks.

| Hemi | Location | ROI | Sensory bias | N | MNI coordinates | | Area (mm$^2$) | |
|---|---|---|---|---|---|---|---|---|
| | | | | | Mean | SD | Mean | SD |
| LH | Lateral | sPCS | Visual | 14 | (-30, -5, 50) | 7, 5, 4 | 284 | 264 |
| | | tgPCS | Auditory | 15 | (-43, -3, 45) | 7, 5, 2 | 293 | 182 |
| | | iPCS | Visual | 15 | (-43, 3, 31) | 6, 5, 5 | 257 | 223 |
| | | cIFS/G | Auditory | 14 | (-45, 13, 20) | 5, 7, 5 | 358 | 206 |
| | | aCO | Auditory | 14 | (-52, 8, 10) | 3, 5, 6 | 303 | 185 |
| | | midIFS | Visual | 14 | (-39, 26, 17) | 6, 7, 6 | 170 | 148 |
| | | FO | Auditory | 15 | (-40, 30, -1) | 5, 5, 6 | 316 | 215 |
| | Medial | cmSFG | Auditory | 14 | (-7, 0, 62) | 2, 6, 4 | 172 | 143 |
| RH | Lateral | sPCS | Visual | 15 | (34, -3, 50) | 7, 4, 4 | 505 | 366 |
| | | tgPCS | Auditory | 15 | (47, -1, 43) | 5, 4, 5 | 277 | 198 |
| | | iPCS | Visual | 15 | (44, 6, 29) | 5, 5, 4 | 353 | 180 |
| | | cIFS/G | Auditory | 15 | (47, 16, 20) | 5, 6, 6 | 403 | 226 |
| | | aCO | Auditory | 14 | (52, 7, 6) | 3, 6, 4 | 223 | 198 |
| | | midIFS | Visual | 14 | (45, 30, 15) | 4, 8, 8 | 203 | 131 |
| | | FO | Auditory | 15 | (45, 28, -3) | 4, 5, 5 | 260 | 219 |
| | Medial | cmSFG | Auditory | 14 | (7, 3, 62) | 2, 6, 3 | 154 | 130 |

Supplementary Table 1: Visual-biased and auditory-biased frontal lobe ROIs defined in Noyce et al. 2022. The MNI coordinates of the centroids and the ROI sizes are reported. These ROIs were defined within individual subjects by the contrast of 2-back visual WM vs. 2-back auditory WM. The visual stimuli were faces (male or female within a block) and the auditory stimuli were animal sounds (dog or cat sounds within a block). These stimuli were chosen to minimize the ability of subjects to assign a semantic label to stimuli and thus promote sensory-based WM.

| Sensory bias | ROI | ROI Name | ID Rate (%) | MNI Coordinates, by hemisphere | | Area (mm²) | |
|---|---|---|---|---|---|---|---|
| | | | | Mean | SD | Mean | SD |
| Auditory | cmSFG | Caudomedial superior frontal gyrus | 80 | (-8, 6, 60) | 1, 5, 3 | 292 | 200 |
| | | | 80 | (8, 15, 58) | 1, 10, 6 | 207 | 98 |
| | tgPCS | Transverse gyrus intersecting precentral sulcus | 100 | (-45, 0, 44) | 4, 7, 3 | 342 | 308 |
| | | | 100 | (49, -1, 43) | 4, 3, 3 | 272 | 152 |
| | cIFS/G | Caudal inferior frontal sulcus/gyrus | 100 | (-45, 17, 20) | 4, 4, 4 | 281 | 236 |
| | | | 90 | (45, 17, 21) | 6, 4, 3 | 288 | 249 |
| | FO | Frontal operculum | 100 | (-43, 29, 0) | 6,4,7 | 475 | 408 |
| | | | 100 | (47, 29, 2) | 5, 2, 6 | 296 | 248 |
| | CO | Central operculum | 100 | (-54, 0, 11) | 4, 8, 6 | 385 | 240 |
| | | | 100 | (57, -4, 14) | 3, 5, 4 | 395 | 284 |
| | PO | Parietal operculum | 90 | (-59, -24, 21) | 2, 6, 3 | 144 | 152 |
| | | | 80 | (55, -24, 24) | 4. 3. 4 | 83 | 67 |
| | pAud | Posterior Auditory | 100 | (-50, -28, 4) | 1, 3, 2 | 4291 | 1143 |
| | | | 100 | (51, -23, 2) | 1, 2, 1 | 4634 | 1393 |
| Visual | sPCS | Superior precentral sulcus | 80 | (-36, -4, 47) | 7, 3, 1 | 208 | 115 |
| | | | 80 | (36, -2, 48) | 8, 4, 2 | 281 | 121 |
| | iPCS | Inferior precentral sulcus | 100 | (-42, 3, 34) | 6, 6, 6 | 143 | 132 |
| | | | 100 | (43, 5, 32) | 5, 6, 5 | 208 | 135 |
| | midIFS | Mid inferior frontal sulcus | 90 | (-40, 21, 23) | 3, 8, 3 | 157 | 91 |
| | | | 70 | (44, 26, 19) | 4, 4, 2 | 231 | 98 |
| | preSMA-V | Pre-Supplementary Motor Area, visual | 100 | (-9, 12, 52) | 1, 8, 7 | 65 | 51 |
| | | | 80 | (9, 14, 50) | 1, 8, 7 | 85 | 49 |
| | pVis | Posterior visual | 100 | (-29, -78, -3) | 3, 4, 3 | 9960 | 1701 |
| | | | 100 | (30, -76, 0) | 2, 4, 4 | 10820 | 2062 |
| Supramodal | preSMA-sm | Pre-supplementary motor area, supramodal | 100 | (-9, 16, 48) | 1, 8, 6 | 178 | 125 |
| | | | 90 | (9, 18, 47) | 1, 9, 7 | 214 | 203 |
| | aIns-sm | Anterior insula, supramodal | 80 | (-30, 24, 1) | 1, 4, 3 | 171 | 82 |
| | | | 100 | (33, 25, -1) | 2, 4, 4 | 177 | 170 |
| | midIFS-sm | Mid inferior frontal sulcus, supramodal | 90 | (-40, 21, 22) | 2, 9, 3 | 346 | 342 |
| | | | 90 | (43, 27, 21) | 3, 5, 4 | 186 | 105 |
| | sPCS-sm | Superior precentral sulcus, supramodal | 80 | (-35, 0, 49) | 6, 6, 1 | 194 | 125 |
| | | | 80 | (34, 3, 50) | 7, 4, 4 | 266 | 256 |
| | iPCS-sm | Inferior precentral sulcus, supramodal | 100 | (-44, 3, 34) | 4, 6, 8 | 289 | 225 |
| | | | 100 | (42, 7, 28) | 6, 4, 4 | 370 | 151 |

Supplementary Table 2: Visual-biased, auditory-biased, and supramodal frontal lobe ROIs defined in Tobyne et al. 2025. The MNI coordinates of the centroids and the ROI sizes are reported. The sensory-biased ROIs were defined within individual subjects by the contrast of 2-back visual WM > (2-back auditory WM + 2-back tactile WM) and by 2-back auditory WM > (2-back visual WM + 2-back tactile WM). Supramodal ROIs were defined by the 3-way intersection of visual WM > visual control, auditory WM > auditory control, tactile WM > tactile control. The visual stimuli were faces (male or female within a block) and the auditory stimuli were animal sounds (dog or cat sounds within a block). Tactile stimuli were raised dot patterns haptically examined with the right index finger. These stimuli were chosen to minimize the ability of subjects to assign a semantic label to stimuli and thus promote sensory-based WM.

| Supramodal | Domain-General | Dice Coefficient | % Supramodal contained in any domain-general ROI |
|---|---|---|---|
| preSMA-sm | 8BM | .33 ± .16 | 81 ± 9% |
| | SCEF | .28 ± .13 | |
| | a32pr | .06 ± .05 | |
| | d32 | .04 ± .04 | |
| ACC-sm | a32pr | .21 ± .10 | 59 ± 23% |
| | d32 | .07 ± .06 | |
| aINS-sm | AVI | .45 ± .13 | 60 ± 14% |
| | FOP5 | .23 ± .10 | |
| sPCS-sm | i6-8 | .10 ± .06 | 13 ± 13% |
| iPCS-sm | 6r | .18 ± .16 | 54 ± 13% |
| | IFJp | .14 ± .10 | |
| | 8C | .11 ± .08 | |
| midIFS/MFG-sm | p9-46v | .41 ± .17 | 73 ± 11% |
| | a9-46v | .09 ± .07 | |
| | 8C | .03 ± .02 | |

Supplementary Table 3. Shows the mean Dice coefficients ± SD for each supramodal and domain-general pair assessed. Only pairs of ROIs in which the supramodal ROI search space overlapped with a domain-general ROI were assessed. The final column indicates the mean percent of supramodal ROI vertices contained in any domain-general ROI ± SD. All supramodal ROIs except for sPCS-sm (13%) had on average at least 50% of their vertices contained in a domain-general ROI. Note that percentages of vertices in the domain-general ROIs also contained in the supramodal ROIs was much lower (mean=28%, one ROI over 50%). This may be due to the fact that we are comparing group average domain-general ROIs derived from a parcellation with parcellation-agnostic individual supramodal ROIs, thus the domain-general ROIs are going to be larger due to the power gained from parcel and participant averaging.

| | **Resting State Connectivity** | | | | | | |
|---|---|---|---|---|---|---|---|
| **Network Connection** | **Original** | | | | **Without Replacement ROIs** | | |
| | **Estimated Marginal Mean** | **SE** | **p-value** | **Cohen's d** | **Estimated Marginal Mean** | **SE** | **p-value** |
| Frontal+Posterior Aud ←→ Frontal+Posterior Aud | .33 | .03 | **p<.001** | 1.10 | .36 | .03 | **p<.0001** |
| Frontal+Posterior Aud ←→ Posterior Vis | -.12 | .02 | **p<.001** | .40 | -.13 | .02 | **p<.0001** |
| Frontal+Posterior Aud ←→ Supramodal | .05 | .02 | **p=.007** | .19 | .05 | .02 | **p=.024** |
| Frontal+Posterior Aud ←→ Frontal Vis | .04 | .02 | **p=.018** | .14 | .03 | .02 | p=.09 |
| Frontal Vis ←→ Posterior Vis | .25 | .04 | **p<.0001** | .84 | .27 | .04 | **p<.0001** |
| Frontal Vis ←→ Supramodal | .17 | .02 | **p<.0001** | .51 | .18 | .03 | **p<.0001** |
| Frontal Vis ←→ Frontal Vis | .50 | .05 | **p<.0001** | 1.67 | .55 | .04 | **p<.0001** |
| Supramodal ←→ Posterior Vis | -.10 | .02 | **p<.0001** | .40 | -.12 | .03 | **p<.0001** |
| Supramodal ←→ Supramodal | .33 | .03 | **p<.0001** | 1.19 | .42 | .04 | **p<.0001** |

Supplementary Table 4. Shows the connectivity estimated marginal means, SEs, and p-values for each network connection analyzed via LME. The "Original" column gives these values for the analysis included in the main manuscript while the "Without Replacement ROIs" column shows these values for the same analysis done without including the ~20% ROIs that did not pass our initial thresholding criteria. Bold p-values indicate significance under FDR correction.

| Network Connection | Resting State Connectivity | | | Task gPPI | | |
|---|---|---|---|---|---|---|
| | **Estimated Marginal Mean** | **SE** | **p-value** | **Estimated Marginal Mean** | **SE** | **p-value** |
| Posterior Aud ←→ Posterior Vis | -.20 | .05 | **p<.0001** | -.10 | .06 | p=.09 |
| Posterior Aud ←→ Frontal Vis | 0.00 | .03 | p=.99 | -.09 | .04 | p=.01 |
| Posterior Aud ←→ Supramodal | .01 | .02 | p=.64 | -.10 | .04 | p=.01 |
| Posterior Aud ←→ Frontal Aud | .29 | .03 | **p<.0001** | -.10 | .03 | **p<.001** |
| Frontal Aud ←→ Posterior Vis | -.10 | .02 | **p<.0001** | .00 | .02 | p=.829 |
| Frontal Aud ←→ Frontal Aud | .33 | .03 | **p<.0001** | -.06 | .02 | p=.006 |
| Frontal Aud ←→ Frontal Vis | .03 | .01 | p=.06 | -.06 | .02 | **p<.01** |
| Frontal Aud←→ Supramodal | .07 | .02 | **p<.01** | -.04 | .02 | p=.04 |
| Posterior Vis ←→ Frontal Vis | .17 | .04 | **p<.0001** | .14 | .05 | **p<.01** |
| Posterior Vis ←→ Supramodal | -.12 | .02 | **p<.0001** | .09 | .03 | **p<.01** |
| Frontal Vis ←→ Frontal Vis | .35 | .04 | **p<.0001** | -.01 | .04 | p=.83 |
| Frontal Vis ←→ Supramodal | .19 | .02 | **p<.0001** | -.03 | .02 | p<.15 |
| Supramodal ←→ Supramodal | .35 | .03 | **p<.0001** | -.02 | .02 | p=.46 |

Supplementary Table 5. Shows the resting state connectivity and task gPPI coefficients estimated marginal means, SEs, and p-values for each network connection analyzed via LME using *a priori* symmetric network clusters. Note that the estimated marginal means for the task gPPI column are derived from subtraction between the visual WM and auditory WM PPI terms, thus positive values indicate visual WM > auditory WM changes in connectivity while negative values indicate auditory WM > visual WM changes in connectivity. Bold p-values indicate significance under FDR correction.

| | Task gPPI | | | | | | |
|---|---|---|---|---|---|---|---|
| **Network Connection** | **Original** | | | | **Without Replacement ROIs** | | |
| | **Estimated Marginal Mean** | **SE** | **p-value** | **Cohen's d** | **Estimated Marginal Mean** | **SE** | **p-value** |
| Frontal+Posterior Aud ←→ Frontal+Posterior Aud | -.07 | .02 | **p<.001** | .26 | -.08 | .02 | **p<.001** |
| Frontal+Posterior Aud ←→ Posterior Vis | -.01 | .02 | p=.58 | - | -.01 | .03 | p=.69 |
| Frontal+Posterior Aud ←→ Supramodal | -.05 | 0.02 | **p=.005** | .20 | -.05 | .02 | **p=.008** |
| Frontal+Posterior Aud ←→ Frontal Vis | -.06 | .02 | **p=.001** | .23 | -.06 | .02 | **p=.002** |
| Frontal Vis ←→ Posterior Vis | .11 | .03 | **p=.009** | .40 | .11 | .04 | **p=.008** |
| Frontal Vis ←→ Supramodal | -.04 | .02 | p=.12 | - | -.04 | .03 | p=.14 |
| Frontal Vis ←→ Frontal Vis | -.03 | .07 | p=.62 | - | .00 | .07 | p=.97 |
| Supramodal ←→ Posterior Vis | .11 | .03 | **p=.001** | .41 | .12 | .03 | **p<.001** |
| Supramodal ←→ Supramodal | -.01 | .02 | p=.50 | - | -.02 | .02 | p=.50 |

Supplementary Table 6. Shows the estimated marginal means, SEs, and p-values for the difference between the visual and auditory WM condition PPI terms in each network connection analyzed via LME. Note that the estimated marginal means are derived from subtraction between the visual WM and auditory WM PPI terms, thus positive values indicate visual WM > auditory WM changes in connectivity while negative values indicate auditory WM > visual WM changes in connectivity. The "Original" column gives these values for the analysis included in the main manuscript while the "Without Replacement ROIs" column shows these values for the same analysis done without including the ~20% ROIs that did not pass our initial thresholding criteria.

| Task gPPI | | | | | | |
|---|---|---|---|---|---|---|
| **Network Connection** | **Original** | | | **Without Fixation in Baseline Conditions** | | |
| | **Estimated Marginal Mean** | **SE** | **p-value** | **Estimated Marginal Mean** | **SE** | **p-value** |
| Frontal+Posterior Aud ←→ Frontal+Posterior Aud | -.07 | .02 | **p<.001** | -.02 | .005 | **p<.001** |
| Frontal+Posterior Aud ←→ Posterior Vis | -.01 | .02 | p=.58 | -.00 | .007 | p=.53 |
| Frontal+Posterior Aud ←→ Supramodal | -.05 | 0.02 | **p=.005** | -.01 | .005 | **p=.006** |
| Frontal+Posterior Aud ←→ Frontal Vis | -.06 | .02 | **p=.001** | -.02 | .005 | **P=.002** |
| Frontal Vis ←→ Posterior Vis | .11 | .03 | **p=.009** | .03 | .01 | **p=.005** |
| Frontal Vis ←→ Supramodal | -.04 | .02 | p=.12 | -.01 | .006 | p=.09 |
| Frontal Vis ←→ Frontal Vis | -.03 | .07 | p=.62 | -.01 | .01 | p=.53 |
| Supramodal ←→ Posterior Vis | .11 | .03 | **p=.001** | .03 | .008 | **p<.001** |
| Supramodal ←→ Supramodal | -.01 | .02 | p=.50 | .00 | .006 | p=.46 |

Supplementary Table 7. Shows the estimated marginal means, SEs, and p-values for the difference between the visual and auditory WM condition PPI terms in each network connection analyzed via LME. Note that the estimated marginal means are derived from subtraction between the visual WM and auditory WM PPI terms, thus positive values indicate visual WM > auditory WM changes in connectivity while negative values indicate auditory WM > visual WM changes in connectivity. The “Original” column gives these values for the analysis included in the main manuscript while the “Without Fixation in Baseline Conditions” column shows these values for the same analysis done without including fixation in the baseline conditions for gPPI analysis.

### Supplementary References

Knief, U., & Forstmeier, W. (2021). Violating the normality assumption may be the lesser of two evils. *Behavior Research Methods*, *53*(6), 2576–2590. https://doi.org/10.3758/s13428-021-01587-5

Noyce, A. L., Lefco, R. W., Brissenden, J. A., Tobyne, S. M., Shinn-Cunningham, B. G., & Somers, D. C. (2022). Extended Frontal Networks for Visual and Auditory Working Memory. *Cerebral Cortex*, *32*(4), 855–869. https://doi.org/10.1093/cercor/bhab249

Schmidt, A. F., & Finan, C. (2018). Linear regression and the normality assumption. *Journal of Clinical Epidemiology*, *98*, 146–151. https://doi.org/10.1016/j.jclinepi.2017.12.006

Tobyne, S. M., Brissenden, J. A., Noyce, A. L., & Somers, D. C. (2025). Combined Auditory, Tactile, and Visual FMRI Reveals Sensory-Biased and Supramodal Working Memory Regions in the Human Frontal Cortex. *Journal of Neuroscience*, *45*(38). https://doi.org/10.1523/JNEUROSCI.0773-25.2025
